# A systematic review of machine learning on clinical MALDI-TOF MS

**DOI:** 10.1101/2025.01.25.634879

**Authors:** Lucía Schmidt-Santiago, Alejandro Guerrero-López, Carlos Sevilla-Salcedo, David Rodríguez-Temporal, Belén Rodríguez-Sánchez, Vanessa Gómez-Verdejo

## Abstract

Bacterial identification, antimicrobial resistance prediction, and strain typification are critical tasks in clinical microbiology, essential for guiding patient treatment and controlling the spread of infectious diseases. While Machine Learning (ML) has shown immense promise in enhancing Matrix-Assisted Laser Desorption/Ionization-Time of Flight Mass Spectrometry (MALDI-TOF MS) applications for these tasks, there is currently no comprehensive review that fully addresses this from a technical ML perspective. To address this gap, we systematically reviewed 115 studies published between 2004 and 2025, focusing on key ML aspects such as data size and balance, pre-processing pipelines, model selection and evaluation, open-source data, and code availability. Our analysis highlights the predominant use of classical ML models like Random Forest and Support Vector Machines, alongside emerging interest in Deep Learning approaches for handling complex, high-dimensional data. Despite significant progress, challenges such as inconsistent pre-processing workflows, reliance on black-box models, limited external validation, and insufficient open-source resources persist, hindering transparency, reproducibility, and broader adoption. This review offers actionable insights to enhance ML-driven bacterial diagnostics, advocating for standardized methodologies, greater transparency, and improved data accessibility. In addition, we provide guidelines on how to approach ML for MALDI-TOF MS analysis, helping researchers navigate key decisions in model development and evaluation.

## Introduction

Identifying and characterizing microorganisms is essential in clinical microbiology to diagnose infections, guide effective antibiotic therapy, and support infection control [1]. Accurate and timely identification, along with determination of antimicrobial resistance (AMR) profiles through Antibiotic Susceptibility Testing (AST), enables clinicians to select appropriate treatments and reduce hospital-acquired infections by up to 30% [2, 3]. However, standard AST methods—such as disk diffusion, broth microdilution, and automated systems—are resource-intensive and slow, often taking 24 to 72 hours [4, 5], which can delay treatment decisions and hinder infection control. Beyond AMR, bacterial strain typification (TYP) is essential for disease management and epidemiological surveillance. TYP is based on molecular techniques such as Polymerase Chain Reaction (PCR), Pulsed-Field Gel Electrophoresis (PFGE), and MultiLocus Sequence Typing (MLST), although the field is increasingly shifting towards Whole-Genome Sequencing (WGS) for its higher resolution. However, WGS remains costly, technically demanding, and often challenging to interpret, thereby limiting its routine use in diagnostic laboratories [6].

The introduction of Matrix-Assisted Laser Desorption/ Ionization-Time of Flight Mass Spectrometry (MALDI-TOF MS) in the late 1980s [7] revolutionized microbiology by enabling rapid, reliable, and cost-effective microorganism identification [8, 9]. By analyzing protein mass spectra, it provides species-level results in under 5 minutes, with a low cost per sample (0.5€/sample) and high-throughput capabilities, making it a standard in clinical laboratories [10].

The technique involves applying an organic matrix to the sample, enabling laser-induced ionization. The resulting ions are separated by their mass-to-charge (m/z) ratio in a Time-of-Flight analyzer, generating a high-dimensional spectrum that serves as a molecular fingerprint for each microorganism [11–13].

Recent advances in Machine Learning (ML) have shown great potential to automate microbial identification (ID) and antimicrobial resistance (AMR) prediction from MALDI-TOF MS data [14–23]. Algorithms such as Support Vector Machines, Random Forests, and Neural Networks can extract complex patterns from spectra, reducing workload and enabling faster, more accurate diagnostics [24–26]. These tools also offer promising support for strain characterization and outbreak management [27–31].

However, challenges remain, particularly the need for large, high-quality datasets, standardized pre-processing, and solutions to class imbalance—especially in AMR and strain typing (TYP). Emerging approaches such as generative Artificial Intelligence (AI) and advanced Deep Learning (DL) models can help address these issues through data augmentation [32, 33], enhanced interpretability [34], and broader clinical applications. Initiatives like DRIAMS and MARISMa databases [21, 35, 36] highlight the importance of open-access spectral repositories to accelerate progress in the field.

While several reviews have highlighted the promise of ML in microbial diagnostics, most lack a strong technical perspective on model development and validation. For instance, focused on bacterial characterization and AMR prediction up to 2020, emphasizing species and antibiotic coverage but offering only limited discussion of ML workflows. [37] analyzed AMR prediction using ML between 2016 and 2024, but excluded ID and TYP tasks and overlooked key aspects such as data pre-processing, imbalance handling, and hyperparameter tuning. Similarly, [38] adopted a broader scope on AI in antimicrobial stewardship beyond MALDI-TOF MS. Although valuable for microbiologists, these reviews fall short in addressing reproducibility, transparency, and ML-specific design considerations.

To bridge this gap, we present a systematic review of ML applications to MALDI-TOF MS for bacterial ID, AMR prediction, and strain TYP, covering studies from 2004 to 2025. We analyze the complete ML pipeline—from data preparation to clinical validation—with the goal of offering practical guidance for researchers in both microbiology and data science. While referencing [5] for pre-2020 studies, we apply stricter inclusion criteria, focusing exclusively on human bacterial pathogens and ensuring methodological rigor. Specifically, we examine:

- Trends in task scope, data size, instrumentation, and bacterial/antibiotic coverage.
- Exploratory data analysis, pre-processing, feature selection, and data imbalance handling, highlighting the lack of homogenous pre-processing pipelines.
- Model selection, highlighting black-box reliance and its impact on transparency, interpretability, and reproducibility.
- Evaluation metrics, Cross-Validation (CV), and external validation, identifying gaps in standardization.
- Open-source challenges in dataset and software accessibility, proposing strategies for greater transparency and reliability.
- Limitations of ML in MALDI-TOF MS and future directions for clinical integration.

In doing so, we propose best practices to promote reproducible, interpretable, and clinically applicable ML workflows. This review serves as a methodological roadmap for the next generation of AI-enhanced microbiology. Figure 1 summarizes the structure of the paper.

**Fig. 1.**
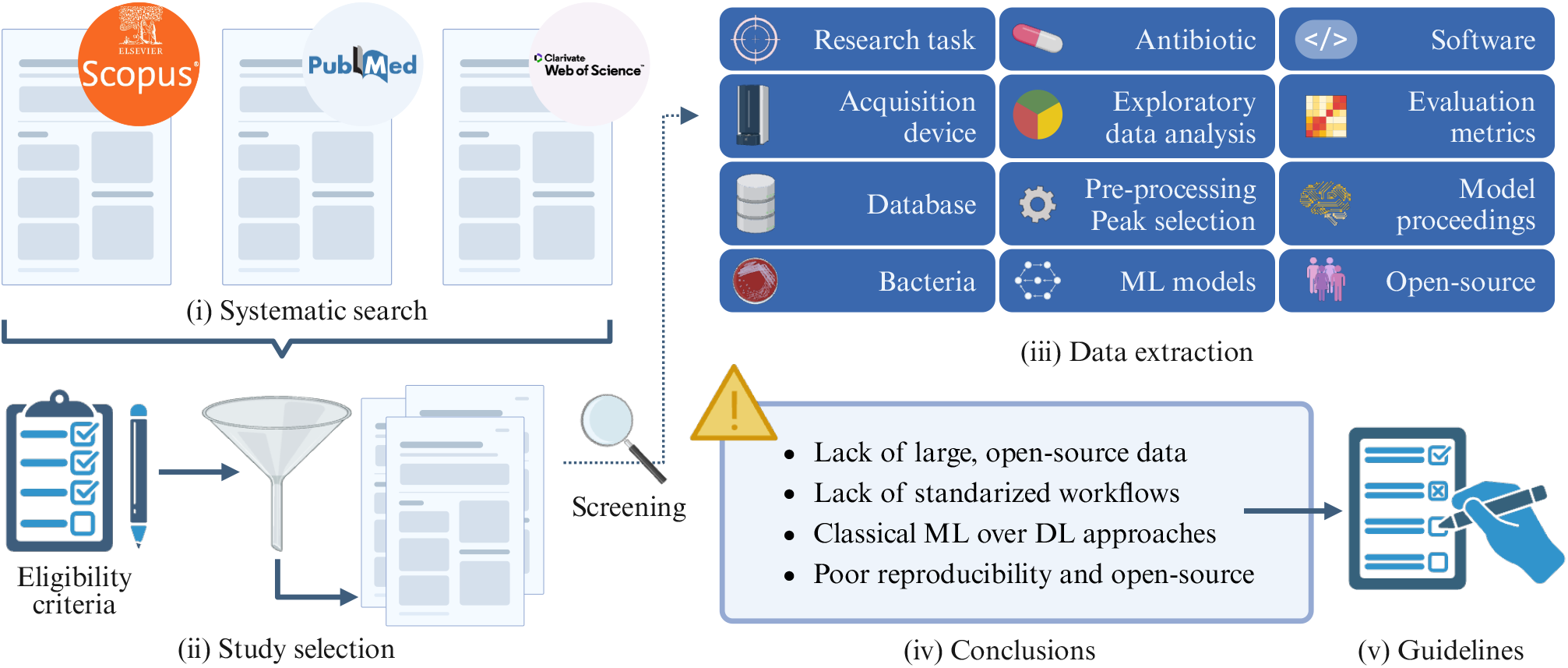
Graphical overview of the systematic review process on Machine Learning (ML) and Deep Learning (DL) applications to MALDI-TOF mass spectrometry data. The workflow includes (i) systematic search in Scopus, PubMed, and Web of Science databases, (ii) study selection based on eligibility criteria, (iii) data extraction including information about research task, acquisition device, bacterial species, databases, antibiotics, preprocessing steps, ML models, evaluation metrics, software and open-source availability, (iv) summary of key limitations identified in the literature, and (v) definition of guidelines for future research.

## Methods

### Search Strategy

A comprehensive literature search was conducted using PubMed/Medline, Scopus, and Web of Science, covering studies published from January 1^st^, 2020, to December 31^st^, 2025. The search aimed to identify studies that applied ML techniques to the analysis of MALDI-TOF MS data. The search included related terms^1^ applied to titles, abstracts, and keywords when available. After retrieving 2,616 documents and incorporating 36 earlier studies from Weis et al. [5], the study selection process was further refined by applying exclusion criteria to ensure the relevance and quality of the studies. The following exclusion criteria were applied:

- **Duplicate records** across repositories (*PubMed/Medline, Scopus*, and *Web of Science*) were removed during the initial screening phase.
- **Irrelevant type records**, such as review articles, letters, conference proceedings, or non-full-text articles in English, were excluded.
- **Off-topic studies** unrelated to MALDI-TOF MS and AI-based methods, such as research focused on genetics, proteomics, cancer, or other unrelated fields, were excluded.
- Articles focusing on **non-human bacteria**, such as those related to foodborne pathogens, animal studies, viruses, or fungi, were removed.
- **Off-task studies** that did not directly address ID, AMR, or TYP were excluded. Examples include studies assessing risk factors, mortality, morbidity prediction, or case studies.

The PRISMA flow diagram (Figure 2) summarizes the screening process and the final set of included studies. This process led to a final selection of 115 studies published between 2004 and 2025. The overlap with previous reviews such as Weis et al. [5], López-Cortés et al. [39], and Pennisi et al. [38] was limited due to our stricter, ML-specific inclusion criteria.

**Fig. 2.**
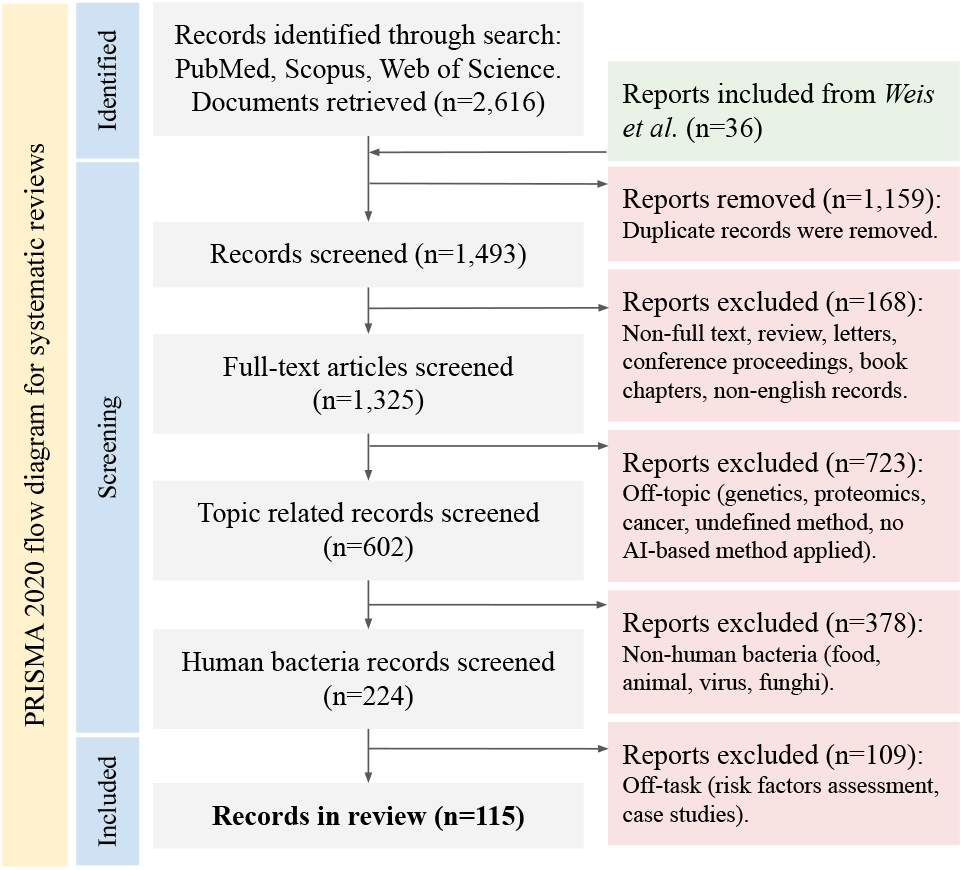
PRISMA 2020 flow diagram illustrating the literature search and screening process. The diagram shows the number of records identified from PubMed, Scopus, and Web of Science databases, and the records included from [5] (n = 36). It details the number of records screened and excluded at each stage based on eligibility criteria, including removal of duplicates, off-topic studies, non-human bacteria, and reports without AI-based methods, resulting in 115 studies.

### Data extraction

To ensure a comprehensive review, we extracted from each study: (i) **publication details**; (ii) the **key research task** (ID, AMR, or TYP); (iii) **bacteria** analyzed; (iv) **antibiotics under study** for AMR tasks; and (v) **data description**, including sample size and spectra details. For computational methodologies, we analyzed (vi) **dataset details and feature importances**; (vii) **pre-processing** and (viii) **feature selection** strategies; (ix) applied **algorithms**; and (x) **Evaluation metrics**. Under **Model proceedings**, we detailed (xi) **Hyperparameter tuning** methods and (xii) the **Evaluation strategy**, including cross-validation and OOD testing. Finally, (xiii) we assessed **Open-source data and code** availability to evaluate transparency and reproducibility. This systematic approach ensured consistency in the literature analysis and facilitated robust comparative analysis across all included studies. Tables 1, 2, and 3 summarize the extracted data.

**Table 1.**
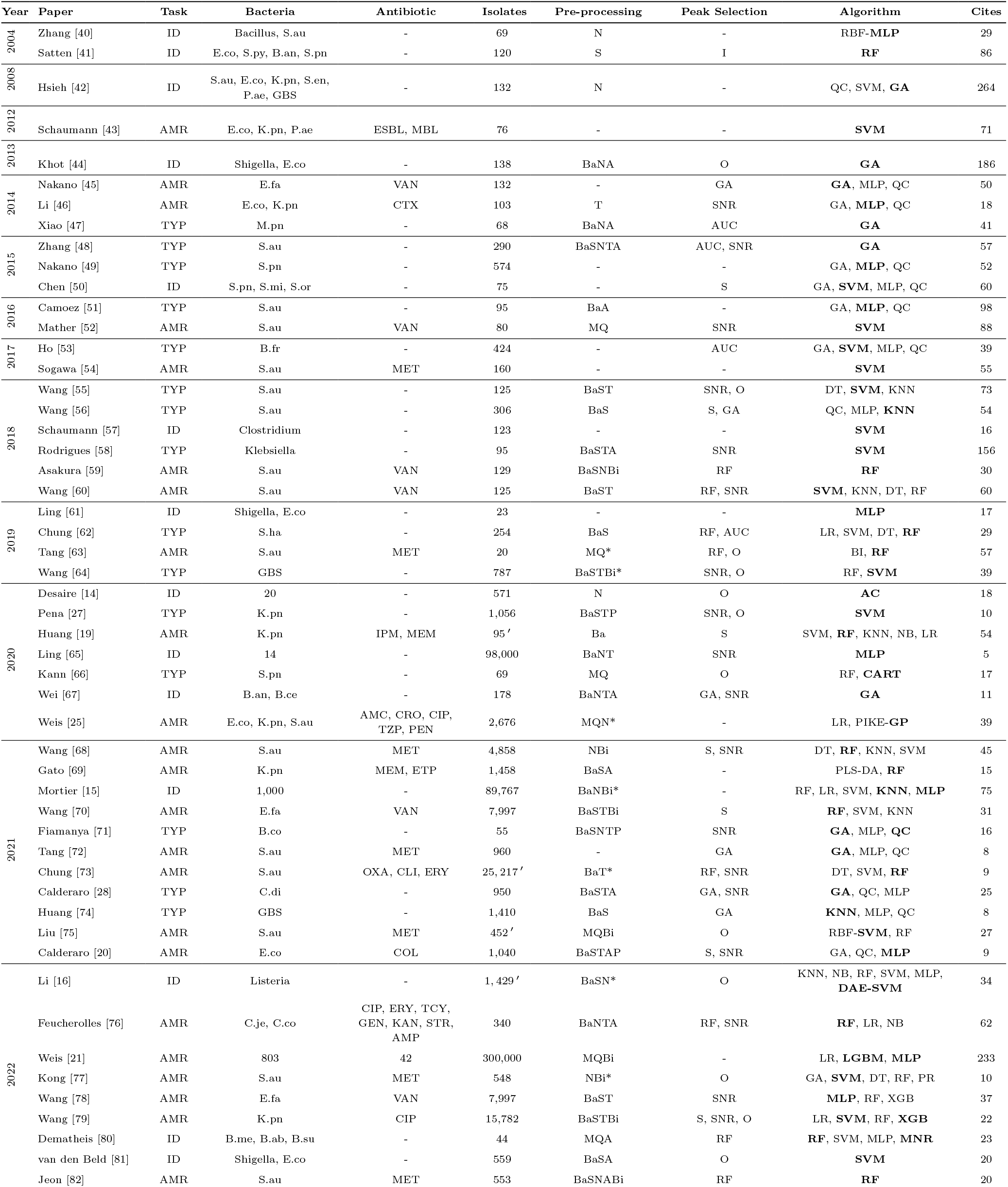
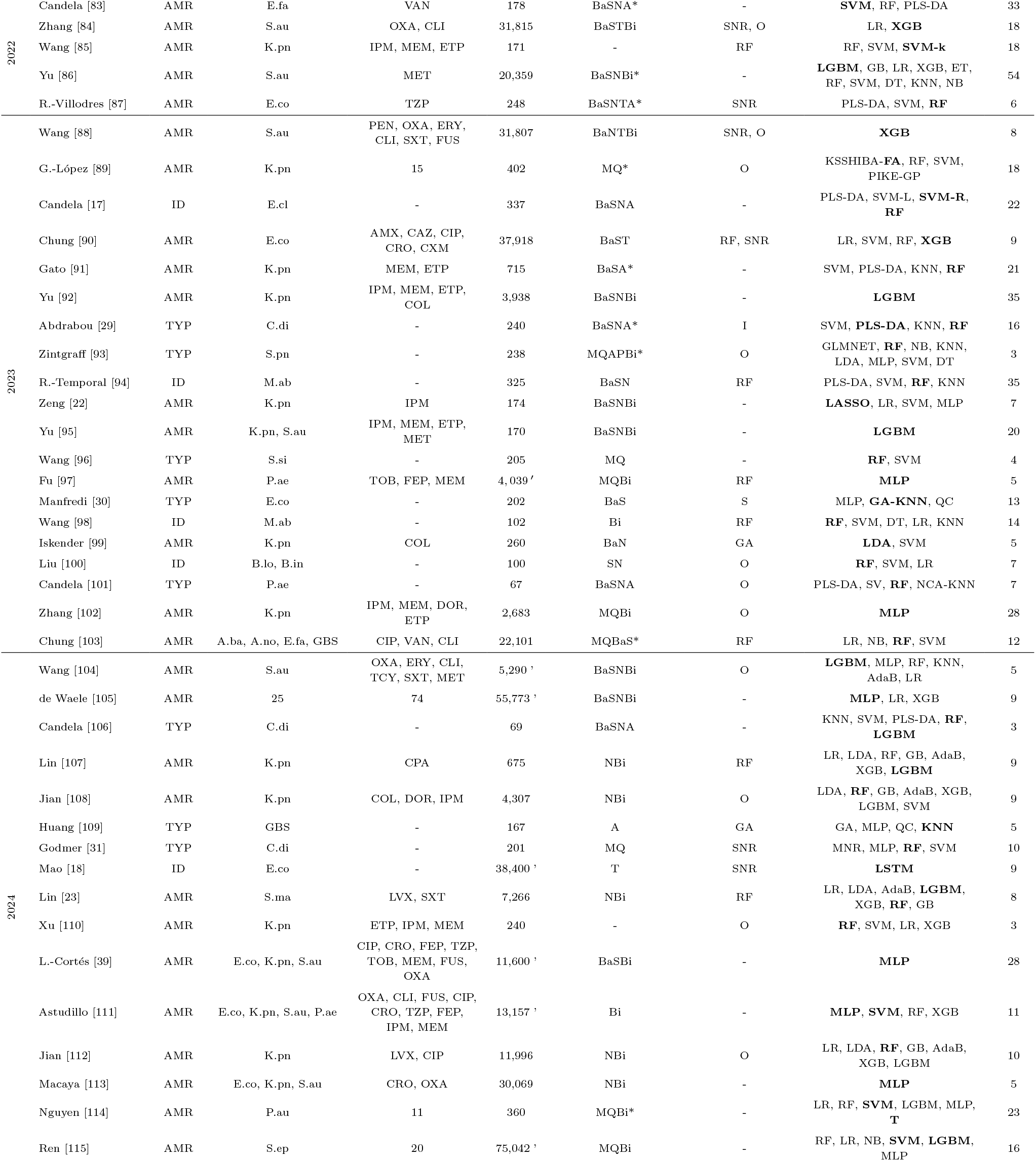
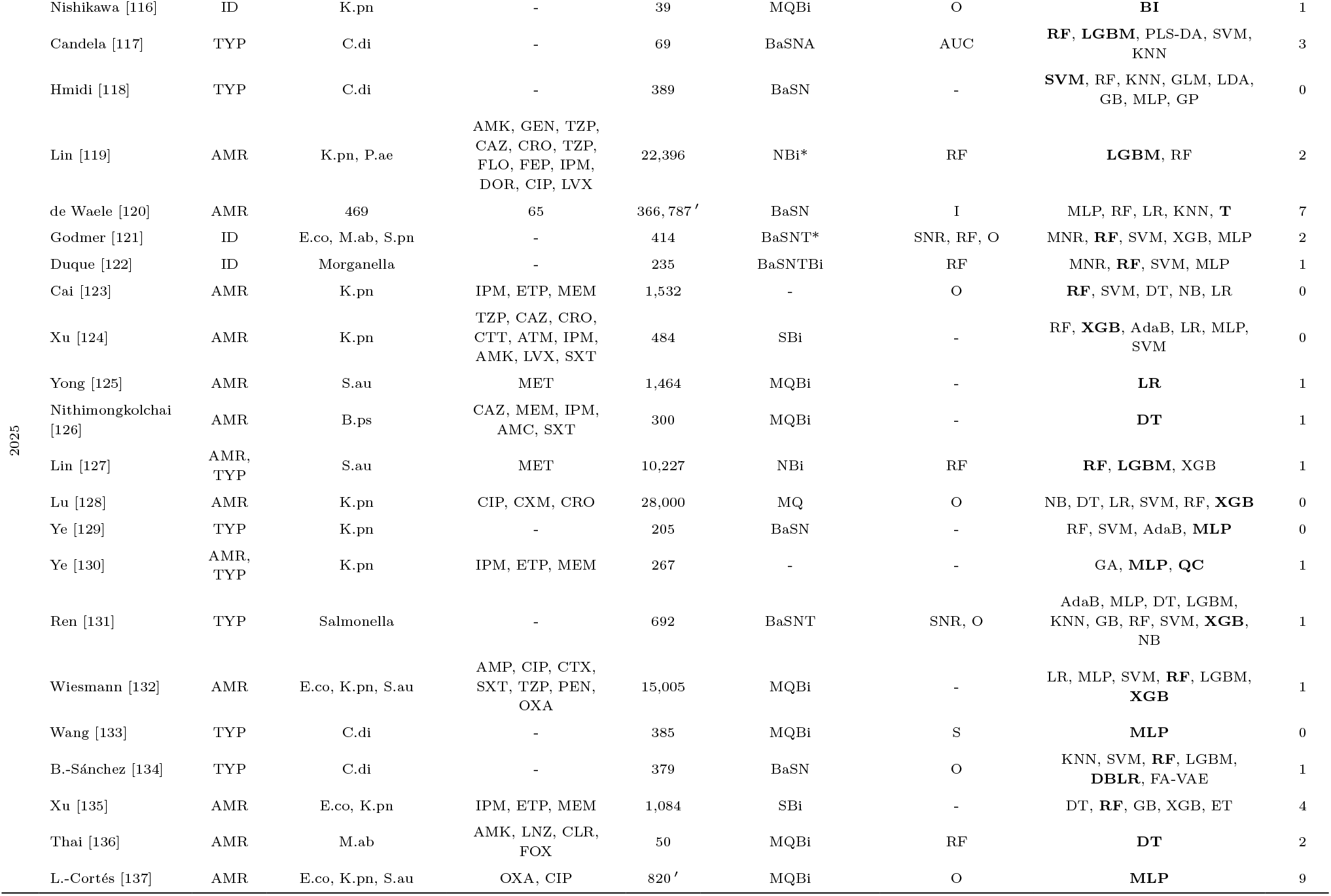
Overview of included studies (2004–2025) categorized by year of publication. The table summarizes 93 studies analyzing MALDI-TOF MS data with Machine Learning (ML) and Deep Learning (DL) approaches. A single quotation mark (‘) in the *Isolates* column indicates spectra; otherwise, it refers to bacterial isolates or samples. Algorithms in bold represent the primary models used or those achieving the best performance. An asterisk (*) in the *Pre-processing* column indicates steps subjected to ablation studies. In the *Cites* column, citation counts obtained from Google Scholar and corresponding to the number of citations recorded as of January 26, 2026, are given. Notation regarding bacterial species, antibiotics, pre-processing steps, and algorithms is explained in 2 and 3.

| Year | Paper | Task | Bacteria | Antibiotic | Isolates | Pre-processing | Peak Selection | Algorithm | Cites |
| --- | --- | --- | --- | --- | --- | --- | --- | --- | --- |
| 2004 | Zhang [40] | ID | Bacillus, S.au | - | 69 | N | - | RBF-MLP | 29 |
|  | Satten [41] | ID | E.co, S.py, B.an, S.pn | - | 120 | S | I | **RF** | 86 |
| 2008 | Hsieh [42] | ID | S.au, E.co, K.pn, S.en, P.ae, GBS | - | 132 | N | - | QC, SVM, **GA** | 264 |
| 2012 | Schaumann [43] | AMR | E.co, K.pn, P.ae | ESBL, MBL | 76 | - | - | **SVM** | 71 |
| 2013 | Khot [44] | ID | Shigella, E.co | - | 138 | BaNA | O | **GA** | 186 |
| 2014 | Nakano [45] | AMR | E.fa | VAN | 132 | - | GA | **GA**, MLP, QC | 50 |
|  | Li [46] | AMR | E.co, K.pn | CTX | 103 | T | SNR | GA, **MLP**, QC | 18 |
|  | Xiao [47] | TYP | M.pn | - | 68 | BaNA | AUC | **GA** | 41 |
| 2015 | Zhang [48] | TYP | S.au | - | 290 | BaSNTA | AUC, SNR | **GA** | 57 |
|  | Nakano [49] | TYP | S.pn | - | 574 | - | - | GA, **MLP**, QC | 52 |
|  | Chen [50] | ID | S.pn, S.mi, S.or | - | 75 | - | S | GA, **SVM**, MLP, QC | 60 |
| 2016 | Camoez [51] | TYP | S.au | - | 95 | BaA | - | GA, **MLP**, QC | 98 |
|  | Mather [52] | AMR | S.au | VAN | 80 | MQ | SNR | **SVM** | 88 |
| 2017 | Ho [53] | TYP | B.fr | - | 424 | - | AUC | GA, **SVM**, MLP, QC | 39 |
|  | Sogawa [54] | AMR | S.au | MET | 160 | - | - | **SVM** | 55 |
| 2018 | Wang [55] | TYP | S.au | - | 125 | BaST | SNR, O | DT, **SVM**, KNN | 73 |
|  | Wang [56] | TYP | S.au | - | 306 | BaS | S, GA | QC, **MLP**, **KNN** | 54 |
|  | Schaumann [57] | ID | Clostridium | - | 123 | - | - | **SVM** | 16 |
|  | Rodrigues [58] | TYP | Klebsiella | - | 95 | BaSTA | SNR | **SVM** | 156 |
|  | Asakura [59] | AMR | S.au | VAN | 129 | BaSNBi | RF | **RF** | 30 |
|  | Wang [60] | AMR | S.au | VAN | 125 | BaST | RF, SNR | **SVM**, KNN, DT, RF | 60 |
| 2019 | Ling [61] | ID | Shigella, E.co | - | 23 | - | - | **MLP** | 17 |
|  | Chung [62] | TYP | S.ha | - | 254 | BaS | RF, AUC | LR, SVM, DT, **RF** | 29 |
|  | Tang [63] | AMR | S.au | MET | 20 | MQ* | RF, O | BI, **RF** | 57 |
|  | Wang [64] | TYP | GBS | - | 787 | BaSTBi* | SNR, O | RF, **SVM** | 39 |
| 2020 | Desaire [14] | ID | 20 | - | 571 | N | O | **AC** | 18 |
|  | Pena [27] | TYP | K.pn | - | 1,056 | BaSTP | SNR, O | **SVM** | 10 |
|  | Huang [19] | AMR | K.pn | IPM, MEM | 95' | Ba | S | SVM, **RF**, KNN, NB, LR | 54 |
|  | Ling [65] | ID | 14 | - | 98,000 | BaNT | SNR | **MLP** | 5 |
|  | Kann [66] | TYP | S.pn | - | 69 | MQ | O | RF, **CART** | 17 |
|  | Wei [67] | ID | B.an, B.ce | - | 178 | BaNTA | GA, SNR | **GA** | 11 |
|  | Weis [25] | AMR | E.co, K.pn, S.au | AMC, CRO, CIP, TZP, PEN | 2,676 | MQN* | - | LR, PIKE-**GP** | 39 |
| 2021 | Wang [68] | AMR | S.au | MET | 4,858 | NBi | S, SNR | DT, **RF**, KNN, SVM | 45 |
|  | Gato [69] | AMR | K.pn | MEM, ETP | 1,458 | BaSA | - | PLS-DA, **RF** | 15 |
|  | Mortier [15] | ID | 1,000 | - | 89,767 | BaNBi* | - | RF, LR, SVM, **KNN**, **MLP** | 75 |
|  | Wang [70] | AMR | E.fa | VAN | 7,997 | BaSTBi | S | **RF**, SVM, KNN | 31 |
|  | Fiamanya [71] | TYP | B.co | - | 55 | BaSNTP | SNR | **GA**, MLP, **QC** | 16 |
|  | Tang [72] | AMR | S.au | MET | 960 | - | GA | **GA**, MLP, QC | 8 |
|  | Chung [73] | AMR | S.au | OXA, CLI, ERY | 25,217' | BaT* | RF, SNR | DT, SVM, **RF** | 9 |
|  | Calderaro [28] | TYP | C.di | - | 950 | BaSTA | GA, SNR | **GA**, QC, MLP | 25 |
|  | Huang [74] | TYP | GBS | - | 1,410 | BaS | GA | **KNN**, MLP, QC | 8 |
|  | Liu [75] | AMR | S.au | MET | 452' | MQBi | O | RBF-**SVM**, RF | 27 |
|  | Calderaro [20] | AMR | E.co | COL | 1,040 | BaSTAP | S, SNR | GA, QC, **MLP** | 9 |
| 2022 | Li [16] | ID | Listeria | - | 1,429' | BaSN* | O | KNN, NB, RF, SVM, MLP, **DAE-SVM** | 34 |
|  | Feucherolles [76] | AMR | C.je, C.co | CIP, ERY, TCY, GEN, KAN, STR, AMP | 340 | BaNTA | RF, SNR | **RF**, LR, NB | 62 |
|  | Weis [21] | AMR | 803 | 42 | 300,000 | MQBi | - | LR, **LGBM**, **MLP** | 233 |
|  | Kong [77] | AMR | S.au | MET | 548 | NBi* | O | GA, **SVM**, DT, RF, PR | 10 |
|  | Wang [78] | AMR | E.fa | VAN | 7,997 | BaST | SNR | **MLP**, RF, XGB | 37 |
|  | Wang [79] | AMR | K.pn | CIP | 15,782 | BaSTBi | S, SNR, O | LR, **SVM**, RF, **XGB** | 22 |
|  | Dematheis [80] | ID | B.me, B.ab, B.su | - | 44 | MQA | RF | **RF**, SVM, MLP, **MNR** | 23 |
|  | van den Beld [81] | ID | Shigella, E.co | - | 559 | BaSA | O | **SVM** | 20 |
|  | Jeon [82] | AMR | S.au | MET | 553 | BaSNABi | RF | **RF** | 20 |
| 2022 | Candela [83] | AMR | E.fa | VAN | 178 | BaSNA* | - | SVM, RF, PLS-DA | 33 |
|  | Zhang [84] | AMR | S.au | OXA, CLI | 31,815 | BaSTBi | SNR, O | LR, **XGB** | 18 |
|  | Wang [85] | AMR | K.pn | IPM, MEM, ETP | 171 | - | RF | RF, SVM, **SVM-k** | 18 |
|  | Yu [86] | AMR | S.au | MET | 20,359 | BaSNBi* | - | **LGBM**, GB, LR, XGB, ET, RF, SVM, DT, KNN, NB | 54 |
|  | R.-Villodres [87] | AMR | E.co | TZP | 248 | BaSNTA* | SNR | PLS-DA, SVM, **RF** | 6 |
| 2023 | Wang [88] | AMR | S.au | PEN, OXA, ERY, CLI, SXT, FUS | 31,807 | BaNTBi | SNR, O | **XGB** | 8 |
|  | G.-López [89] | AMR | K.pn | 15 | 402 | MQ* | O | KSSHIBA-**FA**, RF, SVM, PIKE-GP | 18 |
|  | Candela [17] | ID | E.cl | - | 337 | BaSNA | - | PLS-DA, SVM-L, **SVM-R, RF** | 22 |
|  | Chung [90] | AMR | E.co | AMX, CAZ, CIP, CRO, CXM | 37,918 | BaST | RF, SNR | LR, SVM, RF, **XGB** | 9 |
|  | Gato [91] | AMR | K.pn | MEM, ETP | 715 | BaSA* | - | SVM, PLS-DA, KNN, **RF** | 21 |
|  | Yu [92] | AMR | K.pn | IPM, MEM, ETP, COL | 3,938 | BaSNBi | - | **LGBM** | 35 |
|  | Abdrabou [29] | TYP | C.di | - | 240 | BaSNA* | I | SVM, **PLS-DA**, KNN, **RF** | 16 |
|  | Zintgraff [93] | TYP | S.pn | - | 238 | MQAPBi* | O | GLMNET, **RF**, NB, KNN, LDA, MLP, SVM, DT | 3 |
|  | R.-Temporal [94] | ID | M.ab | - | 325 | BaSN | RF | PLS-DA, SVM, **RF**, KNN | 35 |
|  | Zeng [22] | AMR | K.pn | IPM | 174 | BaSNBi | - | **LASSO**, LR, SVM, MLP | 7 |
|  | Yu [95] | AMR | K.pn, S.au | IPM, MEM, ETP, MET | 170 | BaSNBi | - | **LGBM** | 20 |
|  | Wang [96] | TYP | S.si | - | 205 | MQ | - | **RF**, SVM | 4 |
|  | Fu [97] | AMR | P.ae | TOB, FEP, MEM | 4,039 ' | MQBi | RF | **MLP** | 5 |
|  | Manfredi [30] | TYP | E.co | - | 202 | BaS | S | MLP, **GA-KNN**, QC | 13 |
|  | Wang [98] | ID | M.ab | - | 102 | Bi | RF | **RF**, SVM, DT, LR, KNN | 14 |
|  | Iskender [99] | AMR | K.pn | COL | 260 | BaN | GA | **LDA**, SVM | 5 |
|  | Liu [100] | ID | B.lo, B.in | - | 100 | SN | O | **RF**, SVM, LR | 7 |
|  | Candela [101] | TYP | P.ae | - | 67 | BaSNA | O | PLS-DA, SV, **RF**, NCA-KNN | 7 |
|  | Zhang [102] | AMR | K.pn | IPM, MEM, DOR, ETP | 2,683 | MQBi | O | **MLP** | 28 |
|  | Chung [103] | AMR | A.ba, A.no, E.fa, GBS | CIP, VAN, CLI | 22,101 | MQBaS* | RF | LR, NB, **RF**, SVM | 12 |
| 2024 | Wang [104] | AMR | S.au | OXA, ERY, CLI, TCY, SXT, MET | 5,290 ' | BaSNBi | O | **LGBM**, MLP, RF, KNN, AdaB, LR | 5 |
|  | de Waele [105] | AMR | 25 | 74 | 55,773 ' | BaSNBi | - | **MLP**, LR, XGB | 9 |
|  | Candela [106] | TYP | C.di | - | 69 | BaSNA | - | KNN, SVM, PLS-DA, **RF**, **LGBM** | 3 |
|  | Lin [107] | AMR | K.pn | CPA | 675 | NBi | RF | LR, LDA, RF, GB, AdaB, XGB, **LGBM** | 9 |
|  | Jian [108] | AMR | K.pn | COL, DOR, IPM | 4,307 | NBi | O | LDA, **RF**, GB, AdaB, XGB, LGBM, SVM | 9 |
|  | Huang [109] | TYP | GBS | - | 167 | A | GA | GA, MLP, QC, **KNN** | 5 |
|  | Godmer [31] | TYP | C.di | - | 201 | MQ | SNR | MNR, MLP, **RF**, SVM | 10 |
|  | Mao [18] | ID | E.co | - | 38,400 ' | T | SNR | **LSTM** | 9 |
|  | Lin [23] | AMR | S.ma | LVX, SXT | 7,266 | NBi | RF | LR, LDA, AdaB, **LGBM**, XGB, **RF**, GB | 8 |
|  | Xu [110] | AMR | K.pn | ETP, IPM, MEM | 240 | - | O | **RF**, SVM, LR, XGB | 3 |
|  | L.-Cortés [39] | AMR | E.co, K.pn, S.au | CIP, CRO, FEP, TZP, TOB, MEM, FUS, OXA | 11,600 ' | BaSBi | - | **MLP** | 28 |
|  | Astudillo [111] | AMR | E.co, K.pn, S.au, P.ae | OXA, CLI, FUS, CIP, CRO, TZP, FEP, IPM, MEM | 13,157 ' | Bi | - | **MLP**, SVM, RF, XGB | 11 |
|  | Jian [112] | AMR | K.pn | LVX, CIP | 11,996 | NBi | O | LR, LDA, **RF**, GB, AdaB, XGB, LGBM | 10 |
|  | Macaya [113] | AMR | E.co, K.pn, S.au | CRO, OXA | 30,069 | NBi | - | **MLP** | 5 |
|  | Nguyen [114] | AMR | P.au | 11 | 360 | MQBi* | - | LR, RF, SVM, LGBM, MLP, **T** | 23 |
|  | Ren [115] | AMR | S.ep | 20 | 75,042 ' | MQBi | - | RF, LR, NB, SVM, **LGBM**, MLP | 16 |
| 2025 | Nishikawa [116] | ID | K.pn | - | 39 | MQBi | O | **BI** | 1 |
|  | Candela [117] | TYP | C.di | - | 69 | BaSNA | AUC | **RF, LGBM**, PLS-DA, SVM, KNN | 3 |
|  | Hmidi [118] | TYP | C.di | - | 389 | BaSN | - | **SVM**, RF, KNN, GLM, LDA, GB, MLP, GP | 0 |
|  | Lin [119] | AMR | K.pn, P.ae | AMK, GEN, TZP, CAZ, CRO, TZP, FLO, FEP, IPM, DOR, CIP, LVX | 22,396 | NBi* | RF | **LGBM**, RF | 2 |
|  | de Waele [120] | AMR | 469 | 65 | 366, 787 | BaSN | I | MLP, RF, LR, KNN, **T** | 7 |
|  | Godmer [121] | ID | E.co, M.ab, S.pn | - | 414 | BaSNT* | SNR, RF, O | MNR, **RF**, SVM, XGB, MLP | 2 |
|  | Duque [122] | ID | Morganella | - | 235 | BaSNTBi | RF | MNR, **RF**, SVM, MLP | 1 |
|  | Cai [123] | AMR | K.pn | IPM, ETP, MEM | 1,532 | - | O | **RF**, SVM, DT, NB, LR | 0 |
|  | Xu [124] | AMR | K.pn | TZP, CAZ, CRO, CTT, ATM, IPM, AMK, LVX, SXT | 484 | SBi | - | RF, **XGB**, AdaB, LR, MLP, SVM | 0 |
|  | Yong [125] | AMR | S.au | MET | 1,464 | MQBi | - | **LR** | 1 |
|  | Nithimongkolchai [126] | AMR | B.ps | CAZ, MEM, IPM, AMC, SXT | 300 | MQBi | - | **DT** | 1 |
|  | Lin [127] | AMR, TYP | S.au | MET | 10,227 | NBi | RF | **RF, LGBM**, XGB | 1 |
|  | Lu [128] | AMR | K.pn | CIP, CXM, CRO | 28,000 | MQ | O | NB, DT, LR, SVM, RF, **XGB** | 0 |
|  | Ye [129] | TYP | K.pn | - | 205 | BaSN | - | RF, SVM, AdaB, **MLP** | 0 |
|  | Ye [130] | AMR, TYP | K.pn | IPM, ETP, MEM | 267 | - | - | GA, **MLP**, **QC** | 1 |
|  | Ren [131] | TYP | Salmonella | - | 692 | BaSNT | SNR, O | AdaB, MLP, DT, LGBM, KNN, GB, RF, SVM, **XGB**, NB | 1 |
|  | Wiesmann [132] | AMR | E.co, K.pn, S.au | AMP, CIP, CTX, SXT, TZP, PEN, OXA | 15,005 | MQBi | - | LR, MLP, SVM, **RF**, LGBM, **XGB** | 1 |
|  | Wang [133] | TYP | C.di | - | 385 | MQBi | S | **MLP** | 0 |
|  | B.-Sánchez [134] | TYP | C.di | - | 379 | BaSN | O | KNN, SVM, **RF**, LGBM, **DBLR**, FA-VAE | 1 |
|  | Xu [135] | AMR | E.co, K.pn | IPM, ETP, MEM | 1,084 | SBi | - | DT, **RF**, GB, XGB, ET | 4 |
|  | Thai [136] | AMR | M.ab | AMK, LNZ, CLR, FOX | 50 | MQBi | RF | **DT** | 2 |
|  | L.-Cortés [137] | AMR | E.co, K.pn, S.au | OXA, CIP | 820 | MQBi | O | **MLP** | 9 |

**Table 2.**
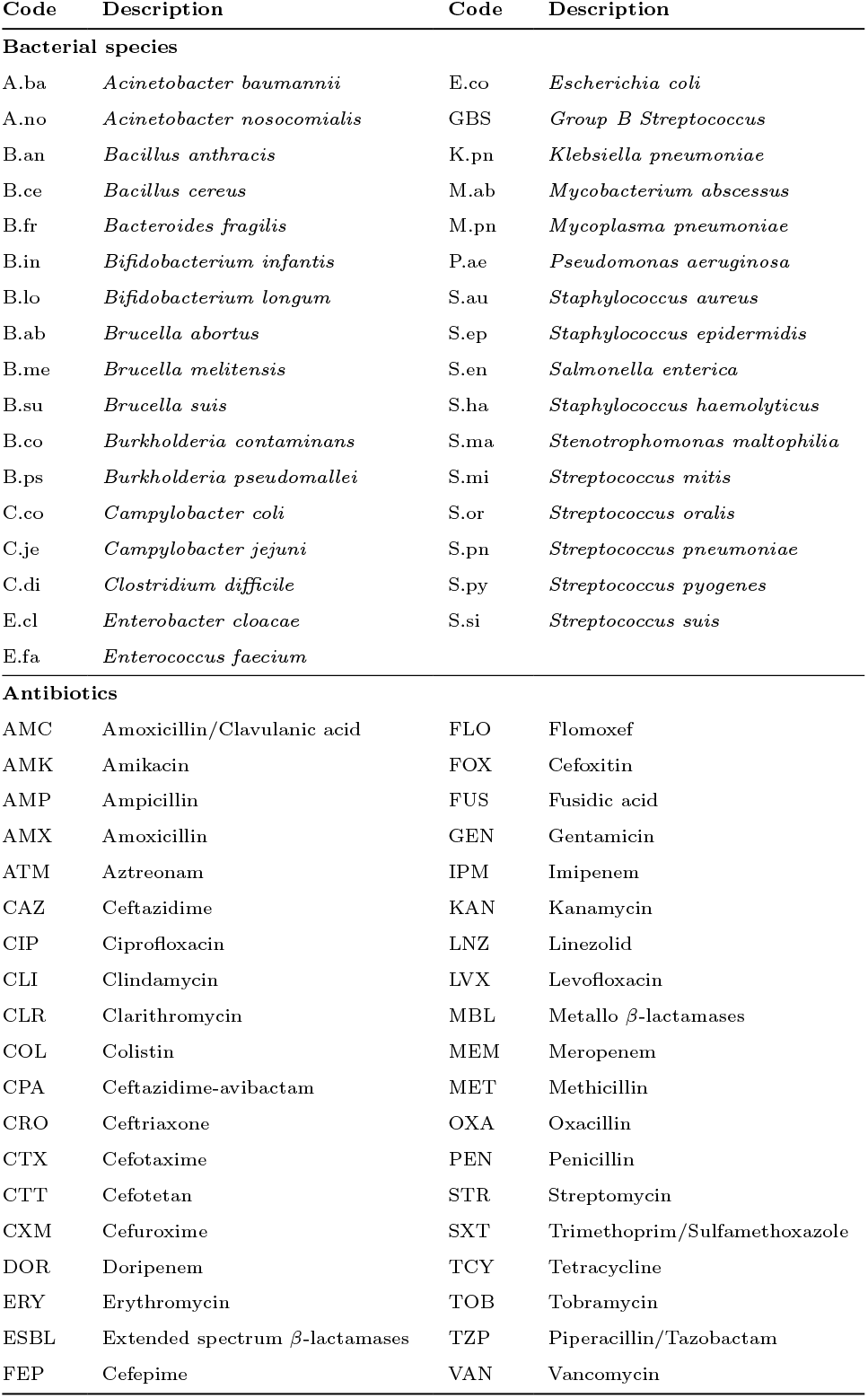
List of codes and abbreviations used for bacterial species and antibiotics.

| Code | Description | Code | Description |
| --- | --- | --- | --- |
| **Bacterial species** |  |  |  |
| A.ba | *Acinetobacter baumannii* | E.co | *Escherichia coli* |
| A.no | *Acinetobacter nosocomialis* | GBS | *Group B Streptococcus* |
| B.an | *Bacillus anthracis* | K.pn | *Klebsiella pneumoniae* |
| B.ce | *Bacillus cereus* | M.ab | *Mycobacterium abscessus* |
| B.fr | *Bacteroides fragilis* | M.pn | *Mycoplasma pneumoniae* |
| B.in | *Bifidobacterium infantis* | P.ae | *Pseudomonas aeruginosa* |
| B.lo | *Bifidobacterium longum* | S.au | *Staphylococcus aureus* |
| B.ab | *Brucella abortus* | S.ep | *Staphylococcus epidermidis* |
| B.me | *Brucella melitensis* | S.en | *Salmonella enterica* |
| B.su | *Brucella suis* | S.ha | *Staphylococcus haemolyticus* |
| B.co | *Burkholderia contaminans* | S.ma | *Stenotrophomonas maltophilia* |
| B.ps | *Burkholderia pseudomallei* | S.mi | *Streptococcus mitis* |
| C.co | *Campylobacter coli* | S.or | *Streptococcus oralis* |
| C.je | *Campylobacter jejuni* | S.pn | *Streptococcus pneumoniae* |
| C.di | *Clostridium difficile* | S.py | *Streptococcus pyogenes* |
| E.cl | *Enterobacter cloacae* | S.si | *Streptococcus suis* |
| E.fa | *Enterococcus faecium* |  |  |
| **Antibiotics** |  |  |  |
| AMC | Amoxicillin/Clavulanic acid | FLO | Flomoxef |
| AMK | Amikacin | FOX | Cefoxitin |
| AMP | Ampicillin | FUS | Fusidic acid |
| AMX | Amoxicillin | GEN | Gentamicin |
| ATM | Aztreonam | IPM | Imipenem |
| CAZ | Ceftazidime | KAN | Kanamycin |
| CIP | Ciprofloxacin | LNZ | Linezolid |
| CLI | Clindamycin | LVX | Levofloxacin |
| CLR | Clarithromycin | MBL | Metallo β-lactamases |
| COL | Colistin | MEM | Meropenem |
| CPA | Ceftazidime-avibactam | MET | Methicillin |
| CRO | Ceftriaxone | OXA | Oxacillin |
| CTX | Cefotaxime | PEN | Penicillin |
| CTT | Cefotetan | STR | Streptomycin |
| CXM | Cefuroxime | SXT | Trimethoprim/Sulfamethoxazole |
| DOR | Doripenem | TCY | Tetracycline |
| ERY | Erythromycin | TOB | Tobramycin |
| ESBL | Extended spectrum β-lactamases | TZP | Piperacillin/Tazobactam |
| FEP | Cefepime | VAN | Vancomycin |

**Table 3.** List of codes and abbreviations used for pre-processing and feature selection steps, as well as machine learning models.

| Code | Description | Code | Description |
| --- | --- | --- | --- |
| **Pre-processing steps** |  |  |  |
| MQ | MALDIquant[138] | Ba | Baseline subtraction |
| S | Smoothing | N | Normalization |
| T | SNR thresholding | A | Averaging |
| Bi | Binning |  |  |
| **Peak selection techniques** |  |  |  |
| AUC | Area under curve | GA | Genetic algorithm |
| I | Intensity thresholding | RF | Random forest |
| S | Statistical tests | SNR | Signal-to-noise ratio |
| O | Other |  |  |
| **Machine Learning algorithms** |  |  |  |
| AC | Aristotle Classifier | LDA | Linear Discriminant Analysis |
| AdaB | AdaBoost | LGBM | Light GB Machine |
| BI | Binary Discriminant Analysis | LR | Logistic Regression |
| CART | Classification Regression Tree | LSTM | Long Short-Term Memory |
| DAE | Denosing Autoencoder | MLP | Multi-Layer Perceptron |
| DBLR | Dual Bayesian LR | MNR | Multinomial Logistic Regression |
| DT | Decision Tree | NB | Naïve Bayes |
| ET | Extra Trees | PLS-DA | Partial Least Squares Discriminant Analysis |
| FA | Factor Analysis | PR | Polynomial Regression |
| FA-VAE | FA Variational Autoencoder | QC | Quick Classifier |
| GB | Gradient Boosting | RBF | Radial Basis Function |
| GLM | Generalized Linear Model | RF | Random Forest |
| GP | Gaussian Process | SVM | Support Vector Machine |
| KNN | k-Nearest Neighbours | T | Transformer |
| LASSO | Least Absolute Shrinkage and Selection Operator | XGB | Extreme GB |

### Quality assessment

Based on the data extracted from each study, following Section Data extraction, we assess its quality according to the following key aspects:

- **ML methodology:** Clear description of the algorithms used, including model architectures, hyperparameter validation ranges, and training procedures.
- **Model evaluation:** Adequate evaluation metrics and validation strategies to ensure robustness and generalization, such as k-fold inner CV or the use of external datasets for OOD evaluation.
- **Data transparency:** Availability of datasets and/or code to enable reproducibility.
- **Clinical relevance:** The practical application of the findings, particularly in ID and TYP bacterial species and predicting AMR in clinical settings.

Each of these aspects will be explored in detail in the Results section, while broader implications and key challenges will be discussed in the Challenges section.

## Results

This section presents the main findings from the reviewed studies (n=115^2^).

### Key research task

Firstly, in ML it is important to distinguish between multi-class, multi-label, and multi-task problems. Multi-class classification assigns one label from several possible classes (e.g., predicting a single bacterial species). Multi-label classification allows multiple binary labels per sample (e.g., resistance to several antibiotics). Multi-task learning combines multiple prediction objectives—such as jointly performing multi-class ID and multi-label AMR prediction—within a single model.

Only a relatively small proportion of the reviewed studies focus on bacterial species identification (n=22). This limited representation is expected, as MALDI-TOF MS is already a gold-standard technique for ID in routine clinical microbiology, often achieving high accuracy without requiring ML-based methods. Consequently, ML is typically applied in ID tasks only when addressing particularly challenging scenarios, such as differentiating closely related species or genera. Examples include species within the *Morganella* genus [122] or discrimination between phylogenetically similar organisms like *Escherichia coli* and *Shigella* [30, 139]. These studies highlight that, while conventional MALDI-TOF MS workflows perform well in standard settings, ML can provide additional resolution in complex classification problems.

AMR remains the dominant research focus across the literature, accounting for n=64 studies. Most works aim to predict resistance profiles to specific antibiotics, reflecting the strong clinical demand for rapid susceptibility assessment. Among studies addressing multiple AMR profiles for bacterial species is typically perform binary classification by training separate models for each resistance phenotype (n=24), the majority do not adopt multi-label learning strategies [21, 23, 25, 39, 73, 76, 88–90, 92, 97, 103, 104, 108, 112–115, 124, 126, 128, 132, 136, 137]. Only n=5 studies employing genuine multi-label classification frameworks [84, 105, 111, 119, 120]. Notably, only one study simultaneously addresses bacterial ID and AMR prediction within a unified modeling framework [105], underscoring the limited exploration of multi-task approaches. Strain typing (TYP) is addressed in n=31 studies. While historically less prevalent than AMR, a notable increase in TYP-focused publications has been observed in 2025, where such studies account for approximately 36% of the papers published, compared to roughly 25% in earlier periods. This recent growth suggests a renewed interest in bacterial strain characterization, likely driven by the increasing importance of epidemiological surveillance [131], outbreak detection [117, 134], and the need to differentiate strains exhibiting distinct resistance profiles [127, 129].

[utab]

#### Summary

Task distribution across the reviewed studies reflects the clinical maturity of MALDI-TOF MS for bacterial ID, where ML is primarily used for challenging discrimination scenarios rather than routine ID. AMR prediction remains the predominant application, consistent with its direct impact on patient management. In contrast, the recent increase in TYP-focused research indicates growing recognition of the importance of strain-level characterization. Figure 3 shows trends in researched tasks over the years. Despite advances, multi-label and multi-task learning strategies remain underexplored, representing a key opportunity for improving the efficiency and clinical applicability of ML-based MALDI-TOF MS workflows.

**Fig. 3.**
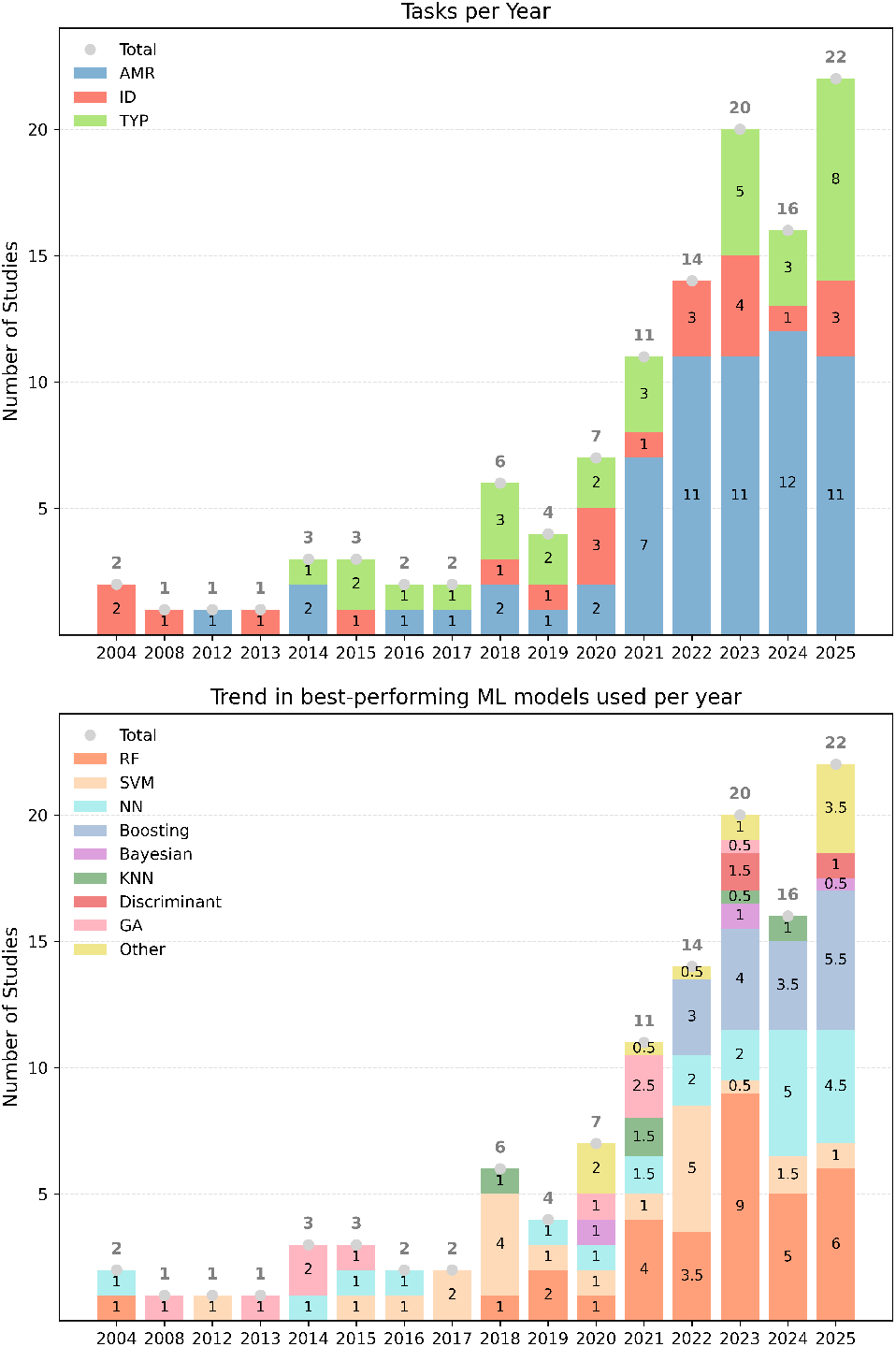
Overview of tasks addressed (top) and best-performing ML models used (bottom) per year across studies. In studies (n=19) reporting two models as equally best-performing (see **bold** in *Algorithm* in Table 1), each model contributes 0.5 to the counts in this visualization.

### Data description

Before applying any ML model, it is essential to understand the dataset, particularly distinguishing between bacterial isolates and mass spectra. Isolates represent cultured strains from clinical or environmental sources, while spectra are their corresponding protein fingerprints obtained through MALDI-TOF MS analysis [140, 141]. One isolate may yield multiple spectra due to technical or biological replicates. However, studies vary in reporting—some specify isolates [75], others only spectra [16], or both [29]—leading to inconsistencies. Since isolates are the most relevant unit for ML, the lack of standardized data reporting complicates study comparisons.

Dataset sizes in the reviewed studies vary significantly, ranging from just 44 isolates [27] to over 300,000 [21, 120]. The majority (n=69) rely on small datasets with fewer than 500 isolates, while n=18 use medium-sized datasets (500–10,000 isolates), and only n=13 incorporate more than 10,000 samples. The median dataset size is 244 isolates and 1,429 spectra.

Data imbalance is a frequent issue, with less than one-third of studies (n=32) working with balanced datasets [18–20, 22, 41, 42, 45, 46, 51, 53, 54, 63, 65, 68, 70, 72–74, 78, 82, 83, 85–87, 91, 93, 98–100, 103, 116, 123], while n=83 report imbalances, particularly in AMR and TYP tasks. Notably, n=10 studies fail to disclose balance status [29, 30, 40, 43, 55, 56, 60, 61, 96, 105]. To address this, n=24 studies apply resampling methods [17, 23, 25, 31, 64, 79, 80, 84, 89, 94, 97, 107, 108, 111– 113, 118, 119, 121, 124, 132, 134, 135, 137], including SMOTE [142] (n=16) [31, 79, 97, 111, 113, 121, 124, 135, 137]. Others (n=14) use imbalance-aware evaluation metrics such as area under the precision-recall curve (AUPRC) or balanced accuracy [16, 21, 25, 31, 39, 76, 89, 97, 101, 102, 104, 111, 113, 115, 120, 124, 125, 128, 133, 134, 137], while only n=4 adjust loss functions [25, 110, 114, 132]. It is concerning that n=35 acknowledge working with imbalanced datasets without applying corrective techniques [14, 15, 27, 28, 44, 47–50, 52, 57– 59, 62, 66, 67, 69, 71, 75, 77, 81, 88, 90, 92, 95, 106, 109, 117, 122, 126, 127, 129–131, 136]. Dataset imbalance is an inherent characteristic of computational microbiology datasets, reflecting the natural distribution of species, strains, and resistance phenotypes. However, transparent reporting of class distributions and explicit consideration of imbalance during model development remain crucial, or otherwise require clear methodological justification.

The reviewed studies employ various MALDI-TOF MS data acquisition devices. The main manufacturer is Bruker Daltonics [143] (n=89), preferred due to its high-throughput capabilities and well-established software tools, such as flexAnalysis and ClinProTools, which facilitate data analysis and integration with ML algorithms. The VITEK MS system by BioMérieux [146] (n=16) lacks the widespread adoption of research seen with Bruker systems, although both provide a full pipeline from measurement to species identification via database lookup. In addition, n=10 studies make use of less common or custom MALDI-TOF instruments, such as Zybio Inc., Applied Biosystems, PerSeptive Biosystems, Absciex, ASTA MicroIDSys, or Voyager DE-STR. Three studies [104, 121, 123] use data acquired from more than one manufacturer.

#### Summary

Establishing systematic data handling and reporting practices—explicitly stating dataset size, biological and technical replicates, and their management—is crucial to prevent bias, misinterpretation, and irreproducibility, ensuring scientific rigour. The lack of medium-large datasets remains a major challenge, though large repositories are beginning to mitigate this issue. Smaller datasets, while easier to handle, often hinder model reliability and generalizability, especially without robust external validation. Dataset imbalance emerges as a pervasive and inherently expected characteristic of computational microbiology data; however, mitigation and evaluation strategies are applied inconsistently, with several studies acknowledging imbalance without corrective measures. Finally, MALDI-TOF MS data acquisition is strongly dominated by Bruker Daltonics systems, while new manufacturers have entered the market in recent years, their adoption remains regionally limited.

### Bacteria

The reviewed studies primarily focus on a few bacterial species that are clinically significant due to their association with hospital-acquired infections and AMR. The most prevalent species studied is *Klebsiella pneumoniae* (n=31), reflecting its critical role in carbapenem-resistant infections, which pose a major challenge in healthcare settings [145]. Following closely, *Staphylococcus aureus* (n=30), which is extensively studied particularly for the detection of methicillin-resistant *S. aureus* (MRSA), a notorious pathogen responsible for numerous healthcare-associated infections [146, 147]. *Escherichia coli* (n=20), known for its role in urinary tract infections and rising resistance to antibiotics such as fluoroquinolones [148] and carbapenems [149], is also a frequent subject of analysis.

The majority of studies (n=88) focus on bacteria from a single genus, such as *Streptococcus* [66, 74, 93, 96]. In contrast, a smaller subset (n=27) examines bacteria across multiple genera, of which only n=8 address more than three genera. This tendency likely stems from the focus of most AMR-related studies on single-species bacterial analysis for susceptibility to one or multiple antibiotics (see Section Antibiotics under study).

#### Summary

Although the diversity of studied bacteria is broad, covering 41 specified species, research efforts remain heavily concentrated on a limited subset of bacterial species, with approximately 60% of studies focusing on *S. aureus, K. pneumoniae*, and *E. coli*. While this distribution is partly justified by their clinical relevance, its concentration introduces a potential research bias, as methodological advances are repeatedly validated on a small number of organisms, which may not capture the full biological and technical variability inherent to MALDI-TOF MS data. Consequently, reported performance levels may overestimate model robustness and generalizability. Since the release of DRIAMS [21, 35], several studies have incorporated a substantially larger number of bacterial species within a single experimental framework. This shift enables evaluation under more heterogeneous and clinically realistic conditions, opening the door to the development of more generalizable models and more complex learning paradigms. Nevertheless, most investigations still rely on single-task and multi-class formulations, limiting the ability of ML systems to fully capture the multi-label nature of clinical microbiology problems.

### Antibiotics under study

The reviewed studies examine a diverse range of antibiotics to predict AMR across various bacterial species, encompassing a total of 39 different antibiotics and 3 antibiotic groups—namely carbapenems, Extended spectrum *β*-lactamases, and Metallo *β*-lactamases. The scope of antibiotics analyzed per study varies significantly; while n=34 studies focus on a single antibiotic or antibiotic-family (e.g., carbapenems), others (n=30) investigate resistance across a broader spectrum. Notably, the largest number of antibiotics examined in a single study reaches 74 [105]. As shown in Table 1, the number of antibiotics analyzed per study as well as the number of studies about AMR has increased over time, underscoring the growing importance of addressing multiresistant bacteria.

Among the antibiotics studied, carbapenems (CP) are the most frequently analyzed (n=25), including *meropenem* (n=15), *imipenem* (n=15), *ertapenem* (n=10), and *doripenem* (n=3). Their critical role in treating infections caused by multidrug-resistant bacteria makes them a primary focus of AMR research. However, the growing threat of CP-resistant bacteria poses a significant public health threat [150] and has led many studies to classify resistance broadly as CP-resistant vs. susceptible (n=14), and others differentiating between specific CP drugs (n=11).

Beyond CP, other commonly studied antibiotics include *methicillin* (n=12), primarily in the context of MRSA detection [54, 63, 68, 72, 75, 77, 82, 86, 95, 104, 125, 127], and *vancomycin* (n=8), widely analyzed for its role in *Enterococcus faecium* and *vancomycin*-resistant *Enterococci* (VRE) [45, 70, 78, 83, 103]. Additionally, *ciprofloxacin* (CIP) (n=12) is frequently studied in *E. coli* and *K. pneumoniae* for urinary tract and systemic infections [25, 39, 79, 90, 111, 112, 119, 128, 132, 137]. *Oxacillin* (OXA) (n=9) is consistently examined for its relevance in detecting *S. aureus* resistance, while *clindamycin* (CLI) (n=6) is another frequently assessed antibiotic. Lastly, *colistin* (COL) (n=4), a last-resort antibiotic, is commonly investigated for resistance in *K. pneumoniae* and other Gram-negative bacteria [20, 92, 99, 108].

However, the inclusion of multiple antibiotics within a single study does not necessarily imply that AMR prediction is performed jointly for all drugs within a unified modeling framework. In most cases, resistance prediction is still addressed through independent binary classification models. Only n=5 studies adopt multi-label classification strategies for AMR prediction. [84] predict *S. aureus* resistance to two antibiotics, [111] perform species-level multi-label classification across nine antibiotics and four bacterial species, [105] investigate three modeling strategies: a single model for all antibiotics and species, individual models per species, and species–antibiotic pair models. [119] analyze resistance patterns for multiple antibiotics across two bacterial species, and [120] employ a recommender system framework to predict resistance profiles across multiple species and antibiotics.

The World Health Organization Bacterial Priority Pathogens List (WHO BPPL) has served as a tool for guiding research and development (R&D) investments, as well as shaping surveillance and control strategies for AMR [151]. The 2024 BPPL update categorizes antibiotic-resistant bacteria into three priority levels: critical, high, and medium. Analyzing how the literature aligns with these categories, it can be observed that for the critical-priority group where it is included CP-resistant *Acinetobacter baumannii* and *Enterobacterales*, such as *K. pneumoniae* and *E. coli*, in cases of CP and third-generation *cephalosporin* resistance (e.g., CTX, CAZ, CPA, CRO), we find n=0, n=14 and n=13, respectively. In the high-priority category, there are studies for CP-resistant *Pseudomonas aeruginosa* (n=4), MRSA (n=12), VRE (n=5), and *fluoroquinolone*-resistant *Salmonella* and *Shigella* species (e.g., CIP, LVX) (n=0), all of which pose increasing threats due to their resistance profiles and prevalence in healthcare settings. While for the medium-priority group, there is no study for *macrolide*-resistant *Streptococcus pneumoniae* and Group A *Streptococci* (e.g., ERY resistance) and *penicillin*-resistant GBS, which represents growing risks, particularly in low- and middle-income countries. These gaps highlight the need for further ML-driven research tailored to high-burden-resistant bacteria, aligning with global AMR mitigation efforts.

#### Summary

In recent years, AMR prediction has become increasingly dominant, drawing research focus away from other tasks, particularly ID. Since 2022, it has been the primary focus of 58% of the reviewed studies. However, AMR predictions are predominantly limited to binary classifications (resistant/susceptible), overlooking distinctions between individual drugs. Despite the growing number of antibiotics analyzed, only three studies employ multilabel classification to predict multiple resistance profiles simultaneously, highlighting a key research gap. While many analyzed antibiotics align with the 2024 WHO BPPL, critical threats like CP-resistant *A. baumannii* remain unaddressed. Expanding multi-label approaches and broadening pathogen coverage could improve AMR prediction and better support global health priorities.

#### Summary

AMR prediction has emerged as the dominant application of ML in MALDI-TOF MS, progressively displacing other tasks such as bacterial identification. However, most studies formulate resistance prediction as independent binary classification problems, a simplification that does not fully capture the multi-drug and multi-mechanism nature of clinical resistance. While this approach facilitates model development, it may limit clinical applicability, where simultaneous resistance profiling is often required. Notably, several high-priority pathogens and resistance scenarios highlighted in the 2024 WHO BPPL, including CP-resistant *A. baumannii*, remain underexplored or absent. Furthermore, modelling strategies better aligned with clinical reality, such as multi-label approaches, remain surprisingly scarce. Expanding beyond binary formulations and improving coverage of clinically relevant resistance threats represent key opportunities for future research.

### Exploratory data analysis and feature importance analysis

Exploratory Data Analysis (EDA) plays a crucial role in understanding the structure, distribution, and variability of the data and helps identify patterns, inconsistencies, and outliers. In fact, it can help us select appropriate data pre-processing and adequate ML algorithms. Visualization is an essential aspect of EDA, as it enables a more intuitive interpretation of complex data structures, making it easier to uncover meaningful relationships and trends.

A total of n=73 studies incorporate some form of EDA, with dimensionality reduction being the most common approach. Principal Component Analysis (PCA) is the most frequently used technique (n=28) [15, 17, 18, 20, 25, 27, 29, 30, 47, 52, 55, 67, 71, 81, 83, 84, 87, 91, 93, 94, 96, 100, 101, 106, 111, 117, 120, 131], followed by t-Distributed Stochastic Neighbor Embedding (t-SNE) (n=7) [17, 89, 90, 98, 114, 128, 131]. While these techniques are often used for dimensionality reduction to 2 or 3 dimensions, their visual representations can also provide valuable insights for data interpretation. Clustering methods are used to group bacterial strains or resistance profiles based on similarity, providing valuable insights into potential subgroups within the dataset. Such techniques include Unsupervised Hierarchical Clustering Analysis (UHCA) (n=17) [17, 27, 42, 57, 63, 64, 66, 70, 80, 81, 91, 93, 94, 96, 101, 116, 117], dendrograms (n=10) [28, 43, 45, 48, 52, 71, 81, 93, 96, 117], and k-means clustering (n=2) [80, 110], aiding in pattern recognition and data exploration.

When working with very high-dimensional data, feature importance analysis becomes essential for understanding how different variables contribute to predictions and overall model performance. Identifying the most influential features helps refine models, improve interpretability, and optimize computational efficiency. Model-agnostic SHapley Additive exPlanations (SHAP) values are frequently used (n=21) [21, 39, 86, 88, 91, 92, 95, 100, 102, 104, 111, 114, 115, 123–125, 131– 133, 135, 137] to interpret the contribution of different features, offering transparency in model decision-making.

#### Summary

Many reviewed studies apply EDA using dimensionality reduction techniques such as PCA and t-SNE, as well as clustering methods including UHCA and dendrograms, primarily to visualize class separability or explore spectral relationships. Feature importance analysis, particularly through SHAP values, is increasingly adopted to enhance interpretability. However, EDA remains inconsistently applied and often superficially interpreted across studies. Given the complexity of MALDI-TOF MS data — including high dimensionality, variability, missing values, and potential outliers — EDA should play a more central role within ML pipelines. Thorough analysis can reveal anomalies, assess distributional properties, guide imputation strategies, and inform critical downstream decisions such as pre-processing, feature selection, and feature engineering. Moreover, systematic EDA contributes directly to model interpretability by enabling more informed ML decisions, reducing black-box behaviour, and supporting explainable modelling strategies. This aspect is particularly critical in medical applications, where transparency, reliability, and clinical applicability strongly depend on understanding both data characteristics and model behaviour. A more structured and systematically reported use of EDA would therefore improve data comprehension, reduce modelling biases, and enhance the robustness and reliability of ML-based findings.

### Pre-processing

Pre-processing is a critical step in MALDI-TOF MS data analysis, directly influencing ML model performance. The principle of “garbage in, garbage out” [152] strongly applies here, emphasizing the need for standardized workflows to ensure high-quality data input. Spectrometry data typically contains noise, such as baseline shifts and fluctuations, which require correction through baseline subtraction and smoothing techniques. Additionally, not all regions of the mass spectrum are informative; peak selection methods help isolate relevant signals (see Section Feature selection). Normalization is essential due to the wide intensity range in spectra, often spanning thousands of units. Furthermore, spectra acquired from different instruments (see Section Data description) exhibit variability, necessitating standardization to ensure robust, generalizable models. Since MALDI-TOF MS data can vary in length, binning techniques are often applied to create fixed-size inputs for ML models. Lastly, technical replicates—multiple spectra from the same sample—must be carefully handled to prevent duplicated data and potential bias. Two widely used pre-processing tools are the open-source R package MALDIquant [138] (n=23, see Table 1), and ClinProTools [153] (n=21) [20, 28, 30, 42, 44–51, 53, 56, 64, 67, 71, 72, 74, 109, 130], a commercial software integrated into Bruker Daltonics’ [143] systems. Both offer extensive pre-processing options, but the lack of standardization in parameter selection and processing order presents a major challenge, hindering reproducibility and comparability across studies. This variability results in diverse pre-processing pipelines, for example, Weis [25] applied normalization, Savitzky-Golay smoothing, baseline correction, Total Ion Current (TIC) normalization, signal-to-noise ratio (SNR) thresholding, warping, and trimming, whereas Dematheis[80] employed spectral cleaning, transformation, smoothing, baseline correction, and normalization.

Despite methodological discrepancies, several pre-processing steps are consistently applied across studies, including baseline subtraction (n=59), smoothing (n=51; e.g., Savitzky-Golay), normalization (n=52; e.g., TIC), SNR thresholding (n=26), spectra averaging (n=24), and binning (n=44). Additional techniques, such as scaling, spectra, and peak alignment, recalibration, binarization, and transformations (e.g., wavelet or helix matrix), are less commonly reported. Table 1 provides a comprehensive overview of the principal steps applied and whether a study includes ablation or comparative studies, indicated by an asterisk (*).

Importantly, only a limited number of studies systematically evaluate how pre-processing choices influence downstream performance. A notable exception is the MSclassifR study by Godmer et al. [121], which performed a large-scale comparative and ablation analysis of pre-processing strategies. Their results indicate that individual choices for smoothing (Wavelet vs. Savitzky–Golay), baseline correction (SNIP vs. TopHat), and normalization (TIC vs. PQN) often yield statistically comparable performance. In contrast, spectral alignment emerged as a decisive factor, with LOWESS-based alignment significantly outperforming alternative approaches. Furthermore, while intensity transformations showed minor differences, the overall pipeline performance proved more robust than any isolated step. This study highlights several practical takeaways: (i) baseline correction, normalization, and alignment are non-negotiable steps; (ii) alignment quality can dominate performance variations; and (iii) excessive focus on marginally different smoothing or calibration methods may offer limited gains relative to pipeline consistency. Readers seeking a deeper understanding of pre-processing effects are encouraged to consult this work directly [121].

#### Summary

The lack of standardization in MALDI-TOF MS pre-processing impacts data quality, model performance, and reproducibility. Studies vary widely in method selection, ordering, and hyperparameter choices, often without clear justification, making replication and optimization difficult. While tools like MALDIquant and ClinProTools provide automated workflows, their customization options introduce inconsistencies that hinder comparability. Comparative evidence indicates that several widely used techniques are largely interchangeable in terms of performance, while spectral alignment may represent a critical determinant of predictive stability. Consequently, baseline correction, normalization, and alignment emerge as essential steps, whereas fine-grained differences between smoothing or transformation strategies appear to play a secondary role relative to pipeline coherence.

### Feature selection

Feature selection, or peak selection (PS) in this context, is a crucial step in handling high-dimensional datasets. In fact, most reviewed studies (n=82) employ PS techniques to enhance model interpretability, improve computational efficiency, and mitigate overfitting. The most common approach involves applying cutoffs, either based on SNR (n=25) or intensity thresholds (n=3) to eliminate minimally informative variables. Additionally, some studies use statistical methods (n=12), such as Pearson Correlation Coefficient, t-tests, or Analysis of Variance (ANOVA), while AUC-based selection (n=4) prioritizes features with the highest discriminative power.

ML-based PS is also widely adopted, with RF-based importance ranking being the most frequently applied technique (n=23). Genetic Algorithms (GA) (n=8) are explored as well, leveraging evolutionary strategies to optimize feature subsets. Moreover, several studies (n=29) utilize alternative PS methods, such as LASSO (n=3) and Bayesian Automatic Relevance Determination (ARD) analysis (n=1). For a comprehensive overview of the PS techniques used in each study, please refer to Table 1.

#### Summary

PS is very helpful for optimizing MALDI-TOF MS-based ML models, improving interpretability, efficiency, and generalization. Advanced ML-driven approaches, such as RF-based importance ranking, Genetic Algorithms, and LASSO, are highly recommended. However, a key unresolved issue is whether MALDI-TOF data should be processed with a fixed input size, as spectra vary in length, and predefined PS may lead to information loss. This challenge is particularly relevant for ML models requiring fixed inputs, though DL techniques, especially Transformer models, offer new possibilities for handling variable-length spectra.

#### Summary

PS plays a central role in MALDI-TOF MS-based ML pipelines, yet the reviewed literature reveals substantial methodological heterogeneity. PS strategies differ fundamentally in how features are evaluated and should therefore be interpreted with caution. Filter methods (e.g., statistical tests, SNR or intensity thresholds), while computationally efficient and easy to implement, assess peaks independently and ignore inter-peak relationships, potentially overlooking biologically meaningful spectral patterns. Wrapper (e.g., forward/backward selection, recursive feature elimination) and embedded (e.g., LASSO, RF importance) methods account for feature dependencies but may reduce interpretability or introduce instability, particularly in small datasets. Consequently, PS design involves an inherent trade-off between performance, interpretability, and biological plausibility — a balance that remains insufficiently discussed. Importantly, emerging modelling strategies challenge the necessity of predefined PS. Approaches operating on variable-length spectra or sparse representations, including transformer-based models, may reduce information loss and better capture spectral structure. Exploring ML pipelines without PS, or combining PS with representation learning techniques, represents a promising direction for improving robustness and clinical applicability.

### Algorithms

A wide range of algorithms (n=34) is applied across the reviewed studies, with most (n=83) comparing multiple models to assess performance. However, algorithm selection is generally driven by dataset size, quality, and computational constraints rather than the specific task (ID, AMR, or TYP), with minimal correlation between model choice and study objective.

Figure 3 illustrates the evolution of best-performing algorithms over time. The increasing use of ensemble models, such as RF and Boosting, along with the emergence of DL approaches—particularly after the release of the DRIAMS database—reflects a shift toward more complex, data-driven methodologies. Despite this, clinical applications often prioritize interpretability. As a result, classical ML models such as RF (n=37) and SVM (n=22) remain the most widely used, owing to their robustness and ability to handle high-dimensional data. RF is particularly favored for its inherent interpretability, which stems from its ensemble structure and feature importance measures. SVMs also offer interpretability when linear kernels are used; however, this advantage diminishes with more complex kernels. Nevertheless, SVMs continue to be popular, especially in settings where the number of features exceeds the number of samples [73, 75, 77, 79, 83, 85, 114, 115].

Boosting methods (n=20), including GB, AdaB [154], LGBM, and XGB [155], also demonstrate strong performance. Less common techniques include genetic algorithms (GA) (n=10), KNN (n=5), discriminant analysis (PLS-DA, LDA) (n=3), and probabilistic approaches (Gaussian processes, Factor Analysis) (n=3). Overall, classical ML models outnumber DL approaches (n=23)—such as Multi-Layer Perceptrons (MLP), Convolutional Neural Networks (CNN), AutoEncoders (AE), Long Short-Term Memory (LSTM), and Transformer models. DL models are advantageous when applied to raw spectra, as they excel at extracting local patterns from mass spectrometry data. However, their effectiveness depends on large datasets [15, 39, 65, 78], with classical ML models often outperforming them when data is limited [118, 121, 124]. Additionally, n=2 studies use two-step models that combine different algorithms to enhance classification [16, 30]. For example, [16] applied a Denoising AE for dimensionality reduction, feeding its latent space into an SVM.

Only n=2 studies [114, 120] employ transformer-based architectures. Nguyen et al. [114] apply transformers within a transfer learning framework, where a pre-trained bacterial identification model is fine-tuned for AMR prediction using a meta-learning strategy. Their approach subsequently integrates five base classifiers (LR, RF, SVM, LGBM, and MLP), leveraging transformer-derived latent embeddings to achieve the highest predictive performance.

More recently, De Waele et al. [120] introduce a transformer architecture specifically designed for MALDI-TOF MS data, combined with a self-supervised pre-training strategy. Their results demonstrate that transformer-based representations achieve state-of-the-art or competitive performance across multiple downstream tasks, including species ID and AMR prediction. Notably, the benefits of this transformer are most pronounced in complex classification settings, such as large-scale multi-class identification and AMR prediction, while performance gains are less evident in simpler tasks. Furthermore, the authors emphasize the critical role of pretraining, showing that transformers trained from scratch exhibit inferior performance, whereas self-supervised pretraining substantially improves generalization.

#### Summary

The reviewed literature highlights a critical methodological observation: increased model complexity does not inherently guarantee improved predictive performance. Classical ML approaches, particularly Random Forest and boosting-based models, consistently demonstrate strong and stable results across a wide range of MALDI-TOF MS applications, indicating high data efficiency in typical small-sample, high-dimensional settings.

Recent studies [120] provide important nuance to this interpretation. While transformer architectures achieve superior performance across multiple downstream tasks, their advantages appear strongly task-dependent. In AMR prediction, transformer-derived representations consistently outperform MLP baselines in AUROC, suggesting improved spectral embedding quality. However, for simpler species ID scenarios, performance gains are less pronounced, with classical methods remaining competitive. Notably, experimental analyses show that the performance advantage of transformer models diminishes as task complexity decreases, reinforcing that architectural sophistication primarily benefits complex classification problems.

Collectively, these findings caution against equating methodological complexity with performance improvement. Instead, coherent ML model design requires careful alignment between task difficulty, dataset characteristics, and modelling strategy. Meaningful comparisons between algorithms are only valid under comparable experimental conditions. Consequently, selecting models based on data regime and problem structure — rather than methodological novelty alone, emerges as a central principle for developing robust and clinically relevant MALDI-TOF MS-based ML systems.

### Evaluation metrics

The reviewed studies employ various evaluation metrics to assess model performance. Accuracy is the most frequently reported metric (n=92) [14, 17–20, 22, 23, 28–31, 40, 41, 43– 57, 59, 61–65, 71–73, 81–87, 90–104, 106–109, 111, 113, 116– 118, 120–124, 128, 129, 133–137], providing a general measure of correctly classified instances. However, given the potential class imbalances in AMR prediction and bacterial ID and TYP tasks, relying solely on accuracy can be misleading. To address this, sensitivity (recall) (n=70) [16, 17, 19–22, 27–31, 42, 45, 46, 48– 51, 56, 57, 62, 64–68, 71–78, 80, 82–87, 90, 92–100, 102–104, 108, 109, 113, 114, 116, 118, 119, 123, 125, 127–131, 133, 136] and specificity (n=64) [16, 17, 19–22, 27–31, 42, 45, 46, 48– 51, 56, 57, 62, 64–68, 71–78, 80, 82–87, 90, 92–100, 102–104, 108, 109, 113, 114, 116, 118, 119, 123–125, 127, 128, 130, 131, 133, 136] are commonly reported, measuring the model’s ability to correctly identify positive and negative cases, respectively. Positive/negative predictive value (n=27)[18, 23, 27, 29, 31, 39, 42, 46, 70, 74, 76, 83, 87, 93, 94, 106–108, 112, 117– 119, 125, 127, 131, 134, 136]/(n=22[23, 27, 29, 31, 39, 42, 46, 70, 74, 76, 83, 87, 93, 107, 108, 112, 118, 119, 125, 127, 131, 136]) assesses the reliability of predictions, while precision (n=8)[18, 31, 65, 84, 98, 116, 120, 129]—which quantifies the proportion of correctly identified positives out of all predicted positives—is less frequently used but remains valuable when false positives are costly. These metrics are particularly critical in AMR tasks, where misclassifying resistant bacteria as susceptible (false negatives) can have serious clinical implications.

F1-score (n=25), which balances precision and recall, is especially useful for imbalanced datasets [16, 23, 65, 66, 76, 83, 87, 91, 92, 102, 104, 107, 108, 111, 112, 114, 118, 119, 124, 127– 129, 131, 133, 137]. AUROC (n=66) is also widely used [15, 17, 21–23, 29, 30, 39, 55–57, 60–62, 65, 68, 70, 73, 75– 79, 82–92, 94–100, 102–108, 110, 112, 114, 115, 117–120, 123– 125, 127–129, 131, 132, 135, 137], offering an aggregate measure of a model’s discriminative ability across different thresholds. To further address class imbalance, the AUPRC (n=17) has gained popularity [15, 21, 25, 29, 39, 76, 88, 91, 102, 104, 114, 115, 125, 128, 132, 133, 137], as it provides a more reliable assessment when the minority class is of primary interest (see Section Data description).

Less frequently reported metrics include confusion matrix analysis (n=6) [18, 22, 59, 66, 80, 99], Kappa (n=8) [31, 69, 82, 93, 118, 121, 122, 125], Youden’s index (n=3) [46, 60, 100], Matthews correlation coefficient (MCC) (n=4) [64, 68, 73, 128], mean error (ME) (n=2) [39, 73], Jaccard index (n=1) [84], Hamming loss (n=1) [111], and G-measure (n=1) [113].

#### Summary

While traditional metrics such as accuracy, sensitivity, and specificity remain dominant, the increasing adoption of AUROC, AUPRC, and F1-score reflects a growing awareness of the challenges posed by imbalanced datasets. The clinical utility of ML performance metrics depends on the specific diagnostic task and its associated risk. In bacterial ID, clinical applicability is often defined by the model’s ability to provide a high-confidence score that distinguishes between species with overlapping spectral profiles. For TYP, the real-world value lies in the sensitivity of the ML model to detect subtle sub-species variations in the proteome that signal outbreaks or hypervirulent lineages. In the case of AMR prediction, spectral classifiers must be evaluated against established regulatory safety benchmarks; for instance, the ISO 20776-2 and FDA standards for AST require that Very Major Errors (VME) remain below 1.5, as failing to detect a resistant proteomic phenotype can lead to treatment failure [156, 157]. Across all these tasks, moving beyond aggregate metrics like accuracy to class-specific performance is essential for ensuring that ML-driven spectral analysis is safe for routine clinical use.

### Model proceedings

#### Hyperparameter tuning

Hyperparameter tuning is crucial for ML model optimization, directly influencing predictive performance and generalizability. However, the extent of tuning varies significantly across the reviewed studies. A substantial number (n=25) do not disclose any details on hyperparameter selection [16, 27, 30, 42, 43, 45, 46, 49, 52–54, 57–59, 61–64, 69, 72, 92, 93, 100, 126, 130], leaving it unclear whether parameters were optimized or left at default values. Additionally, n=40 rely entirely on default settings [18–20, 28, 29, 41, 44, 47, 48, 50, 51, 65– 67, 70, 71, 78, 79, 81, 83, 85, 87, 94–96, 98, 99, 102, 104, 106, 116, 117, 122, 123, 125, 128, 129, 133, 135, 136].

Among studies that perform hyperparameter optimization, CV is the most common approach (n=40) [15, 17, 21–23, 25, 31, 39, 55, 60, 68, 73–77, 80, 84, 86, 88–91, 97, 101, 105, 107– 110, 112, 113, 115, 119, 121, 124, 127, 131, 132, 134]. The CV method optimizes hyperparameters by creating training and validation partitions within the training set, helping to mitigate overfitting and ensure robust model selection. Only n=4 studies apply Bayesian hyperparameter selection [111, 114, 134, 137].

A smaller subset (n=4) employs manual ablation studies [14, 40, 56, 103, 118], though this approach is often inefficient and lacks reproducibility.

#### Evaluation strategy

Evaluating model performance is a crucial step in ML workflows, ensuring that predictive models generalize well beyond their training data. The reviewed studies employ a variety of evaluation strategies, with two studies not specifying their approach (n=2) [61, 126].

Predefined train-test splits are used in n=64 studies [15, 21– 23, 27, 31, 41, 46, 49, 53, 63, 65, 76, 77, 82, 84, 86, 97, 100, 105, 107–109, 111–113, 115, 116, 119–125, 127, 128, 130, 131, 133– 136]. In these cases, a fixed portion of the dataset is allocated for training, while the remaining data is reserved for posterior evaluation. This approach is computationally efficient but may introduce bias if the split is not representative of the full data distribution. N=30 also apply CV on the predefined training set to improve model robustness [14, 18, 20, 29, 39, 42, 51, 54, 66, 67, 69, 70, 74, 75, 79–81, 89, 99, 102, 110, 116, 121– 124, 128, 130, 133, 135].

Outer CV is the most common evaluation method, applied in n=70 studies [16, 17, 28, 30, 43–45, 47, 48, 50, 55– 58, 60, 64, 68, 71–73, 78, 83, 85, 88, 90–92, 94–96, 98, 103, 104, 106, 114, 117, 118, 129, 132, 137]. This technique repeatedly splits the dataset into training and validation folds, reducing overfitting and providing a more reliable estimate of model performance. Within this group, a subset of studies (n=4) employs stratified CV, which ensures class distribution is preserved across folds—a particularly valuable approach for imbalanced datasets [25, 62, 87, 89]. In contrast, n=7 studies rely on Leave-One-Out (LOO) CV for model validation [19, 40, 42, 52, 59, 101, 121]. This method iteratively trains the model on all but one sample, testing it on the excluded instance until every sample has been used for validation. While LOO-CV provides an almost unbiased estimate of model performance, it comes at a high computational cost and suffers from increased variance due to strong correlations between training sets.

Conversely, lower-fold CV methods reduce variance by training on more distinct subsets but may introduce higher bias when training sets are too small. Given this trade-off, LOO-CV is only justified for very small datasets.

OOD validation, or external validation, refers to the process of evaluating an ML model on an independent dataset that was not used during training or outer CV. This step is crucial for assessing a model’s ability to generalize beyond the data on which it was developed, ensuring its robustness when applied to new or unseen distributions. However, despite its importance, OOD is often overlooked or misunderstood. Many studies conflate OOD validation with standard evaluation methods, such as a predefined test split, that assess performance only within the same dataset. Among the reviewed studies, only n=32 actually conduct OOD [15, 17, 21, 39, 42, 58, 64, 68– 71, 73, 78, 84, 86, 88, 90, 91, 93, 94, 100, 103, 111, 114, 115, 120, 121, 123, 128, 131, 132, 134]. The remaining studies rely solely on internal validation.

#### Summary

Model development practices across the reviewed studies reveal several methodological limitations. A substantial proportion of works (n=25) do not disclose hyperparameter configurations, preventing reproducibility and hindering objective assessment of model optimization. When hyperparameters are reported, it remains essential to clarify whether values correspond to default settings or systematic tuning procedures, preferably conducted via cross-validation. Without appropriate tuning, models may operate under suboptimal configurations, potentially underestimating achievable performance. Similarly, studies reporting performance metrics derived from single train-test splits or non-averaged evaluations lack evidence of predictive robustness, as variability across data partitions is not assessed. Reliable ML pipelines require performance estimates aggregated across multiple iterations or validation folds to ensure stability and generalizability. Finally, despite the widespread adoption of internal validation strategies, only n=32 studies conduct true OOD validation. While outer CV mitigates overfitting within a dataset, it primarily evaluates interpolation rather than robustness to distribution shifts commonly encountered in MALDI-TOF MS applications. Given the variability introduced by laboratory protocols, acquisition devices, and population heterogeneity, models lacking external validation remain vulnerable to performance degradation in real-world clinical settings. This imbalance between internal optimization and external robustness highlights a persistent gap between methodological evaluation and clinical deployment requirements.

### Open-source data and code

The availability of open-source data and code is essential for advancing research, promoting transparency, and ensuring reproducibility in ML studies, including sharing software and implementation details. Python is the predominant software (n=47) [15, 18, 21–23, 25, 27, 39, 65, 70, 73, 76–78, 84–86, 88– 90, 92, 95–97, 100, 102, 103, 105, 107, 108, 110–115, 119, 120, 124, 127–129, 131, 133–135, 137] due to its versatility and extensive ML libraries [155, 160, 161]. Additionally, Python is more commonly associated with recent and complex studies, with 58% of publications in the last three years employing this programming language. Bruker’s ClinProTools [143] (n=21) [20, 28, 30, 42, 44–51, 53, 56, 64, 67, 71, 72, 74, 109, 130] leads among manufacturer-specific tools, followed by Bionumerics [162] (n=2) [58, 81] and AMRQuest [163] (n=2) [82, 125]. R [164] (n=22) [14, 31, 52, 59, 60, 62, 63, 66, 68, 70, 75, 79, 80, 93, 98, 116, 118, 121, 122, 126, 132, 136] is favored for its robust statistical analysis capabilities and specialized mass spectrometry tools [138]. Web-based tools like Clover [165] (n=10) [17, 29, 69, 83, 87, 91, 94, 101, 106, 117] offer streamlined clinical workflows but are less flexible. MATLAB [166] (n=6) [40, 43, 55, 57, 61, 99] is used for its matrix handling and visualization, though its closed-source nature limits adoption. Only n=4 studies omit software details, hindering reproducibility[16, 41, 104, 123].

Among the reviewed studies, only a small subset publicly shares their datasets (n=15) and (n=17) share their ML implementation, which is crucial for full reproducibility (see Table 4). Despite the growing availability of MALDI-TOF MS datasets, only two contain data labeled for TYP [93, 96]. The absence of open-access typed data restricts researchers’ ability to develop and validate models for hypervirulence prediction across bacterial strains.

**Table 4.** Summary of studies that provide open-source datasets and/or code. The table lists the available datasets, their sizes, formats (raw or preprocessed, pp), and whether the corresponding code is publicly accessible. Additionally, it includes the software used in each study and relevant repository links.

| Paper | Data | Size | Format | Code | Software | Links |
| --- | --- | --- | --- | --- | --- | --- |
| Asakura [59] | ✓ | N/A | pp | ✓ | C++/R | GitHub |
| Desaire [14] | ✓ | 571 | raw |  |  | UCI |
| Ling [65] |  |  |  | ✓ | Python | GitHub |
| Weis [25] |  |  |  | ✓ | Python | GitHub |
| Weis [21] | ✓ | 132,097 | raw, pp | ✓ | Python | DRYAD, GitHub |
| Wang [78] |  |  |  | ✓ | Python | GitHub |
| Yu [86] | ✓ | 2,456 | pp |  |  | Zenodo |
| Wang [88] | ✓ | 31,807 | pp | ✓ | R | GitHub |
| G.-López [89] | ✓ | 2,000 | raw, pp | ✓ | Python | GitHub |
| Candela [17] | ✓ | 1,566 | raw |  |  | Clover |
| Chung [90] | ✓ | 37,918 | pp |  |  | GitHub |
| Yu [92] | ✓ | 1,853 | pp |  |  | Zenodo |
| Zintgraff [93] | ✓ | 193 | raw | ✓ | R | Mendeley |
| Wang [96] | ✓ | 205 | pp | ✓ | Python | Article |
| de Waele [105] |  |  |  | ✓ | Python | GitHub |
| Mao [18] | ✓ | N/A | pp |  |  | SimTK |
| L.-Cortés [39] |  |  |  | ✓ | Python | GitHub |
| Astudillo [111] |  |  |  | ✓ | Python | GitHub |
| Nguyen [114] | ✓ | 30 | raw | ✓ | Python | GitHub |
| Ren [115] |  |  |  | ✓ | Python | GitHub |
| de Waele [120] |  |  |  | ✓ | Python | GitHub |
| Godmer [121] |  |  |  | ✓ | R | GitHub |
| B.-Sánchez [134] | ✓ | 379 | raw | ✓ | Python | Zenodo, GitHub |
| Thai [136] | ✓ | 50 | raw |  |  | Mendeley |
| RKI dataset [158] | ✓ | 11,038 | raw |  |  | Zenodo |
| MARISMa [36] | ✓ | 202,700 | raw |  |  | Zenodo |
| MS-UMG [159] | ✓ | 77,342 | pp |  |  | Zenodo |
| **Sum:** | **18** |  |  | **17** |  |  |

Among the open datasets, file formats vary considerably, with few studies (n=2) providing raw spectral data (e.g., *BrukerRaw* format). In contrast, others release raw or pre-processed data in the non-original format (*csv, txt, npy*), limiting its usability for other research purposes. As shown in Table 4, only a few datasets retain the original metadata, such as acquisition time or device information, which is crucial for standardization and reproducibility.

Besides the DRIAMS [35] dataset published by [21], other large public MALDI-TOF MS datasets are available to the research community: the RKI [158], MARISMa [36], and MS-UMG [159] datasets. All provide a comprehensive collection of spectra from various bacterial species, enabling large-scale studies on bacterial ID and AMR (except RKI). These repositories have facilitated replicating and validating existing models and spurred new research. A total of n=9 studies use the DRIAMS and/or RKI datasets for their research [39, 97, 104, 105, 111, 113, 115, 120, 132], illustrating the transformative impact of open-access data on advancing the field.

#### Summary

Open-science practices play a fundamental role in ensuring reliability, transparency, and reproducibility in ML-driven microbiology research. Publicly accessible datasets and well-documented code enable independent validation of reported findings, facilitate methodological comparisons, and allow other laboratories to assess model robustness on their own data. In clinical contexts, where variability across instruments, protocols, and populations is unavoidable, reproducibility becomes a prerequisite for trustworthy model deployment. Beyond reproducibility, open data sharing has the potential to substantially accelerate methodological innovation. Large-scale, heterogeneous MALDI-TOF MS repositories incorporating diverse isolates, strain-level variability, resistance phenotypes, and epidemiological heterogeneity would enable the development of more generalizable models and foster emerging research directions, including representation learning, generative AI, and foundational spectral models. Importantly, aggregating data across laboratories would enhance both biological and technical variability, a critical factor for designing ML systems robust to distribution shifts inherent to real-world clinical environments.

### Challenges and future directions

Progress in ML for microbiology critically depends on data and model transparency. The current literature exhibits substantial heterogeneity and incomplete reporting regarding dataset composition, preprocessing pipelines, and model configurations, limiting reproducibility and independent validation. True scientific progress requires clear documentation of data distributions, replicate handling, preprocessing strategies, and hyperparameter settings. Furthermore, emerging microbiological challenges, including closely related species complexes and high-burden resistant pathogens [151], highlight potential gaps between research trends and clinical needs. While certain well-studied problems continue to dominate the literature, other clinically relevant pathogens remain comparatively underexplored. Aligning methodological research with real-world clinical challenges, supported by standardized, large-scale MALDI-TOF MS repositories, represents an essential direction for future work and could substantially accelerate methodological development while improving clinical relevance [167].

Another major challenge lies in adequately characterizing variability inherent to MALDI-TOF MS data. Spectral variability arising from laboratory protocols, acquisition devices, culture conditions, and temporal drift can substantially influence model behavior. Consequently, thorough exploratory data analysis prior to modelling becomes essential, incorporating both interpretability and variability assessment. Recent studies demonstrate that variability-aware data augmentation strategies can mitigate performance degradation and improve model robustness under heterogeneous acquisition conditions [33]. In parallel, domain adaptation and domain alignment methods represent promising directions for addressing distribution shifts, yet their adoption within MALDI-TOF MS workflows remains unexplored.

Consistency in performance assessment also represents a critical methodological requirement. The reviewed studies reveal substantial variability in reporting practices, including incomplete disclosure of hyperparameter tuning procedures, inconsistent feature selection strategies, and reliance on non-averaged evaluation metrics. Proper ML pipelines require systematic hyperparameter optimization tailored to the specific data environment, alongside rigorous validation strategies. Performance estimates should be aggregated across multiple data folds to ensure robustness. Importantly, clinical ML applications must account for error asymmetry, such as in AMR prediction, where false-negative errors may have disproportionately severe clinical consequences. Consequently, evaluation metrics sensitive to minority-class performance and clinical risk, such as sensitivity, recall, AUPRC, or balanced accuracy, should be prioritized.

Finally, external validation remains one of the most significant barriers to clinical applicability. While internal validation strategies are widely adopted, only a limited number of studies perform true OOD evaluation. Given the variability and distribution shifts inherent to clinical microbiology data, models validated exclusively within a single dataset may exhibit reduced transportability across laboratories or clinical environments. Robust OOD validation is therefore essential for assessing model reliability, stability, and real-world clinical utility [168].

Recent advances in DL and representation learning offer promising directions for MALDI-TOF MS analysis, although DL methods remain limited in the reviewed literature. Observed performance trends indicate that for simpler tasks, such as binary AMR prediction or restricted multi-class classification, increased model complexity does not consistently yield performance gains. Classical ML approaches, particularly RF and boosting-based models, often provide robust and competitive results, highlighting their strong data efficiency. In contrast, representation learning strategies, including multivariate methods such as PLS and transformer architectures leveraging self-supervised pretraining, demonstrate clearer advantages in more complex scenarios by effectively capturing spectral variability and improving generalization. Complementing these developments, recent generative AI approaches enabling conditional spectrum synthesis [169] present a potential solution to data scarcity and class imbalance, offering new opportunities for data augmentation and variability-aware model development.

### Guidelines

After reviewing n=115 SOTA papers, we identified a lack of standardization in ML research on MALDI-TOF MS data. To address this, we propose a set of guidelines to establish a well structured research pipeline that can effectively advance ML applications in this field. A clear definition of the research task is crucial, specifying how data will be utilized and how ML techniques will be applied. Once this is defined, the pipeline should follow key stages: (i) EDA, (ii) spectra pre-processing and peak selection (PS), (iii) model selection, (iv) training, and (v) evaluation, as in Figure 4.

**Fig. 4.**
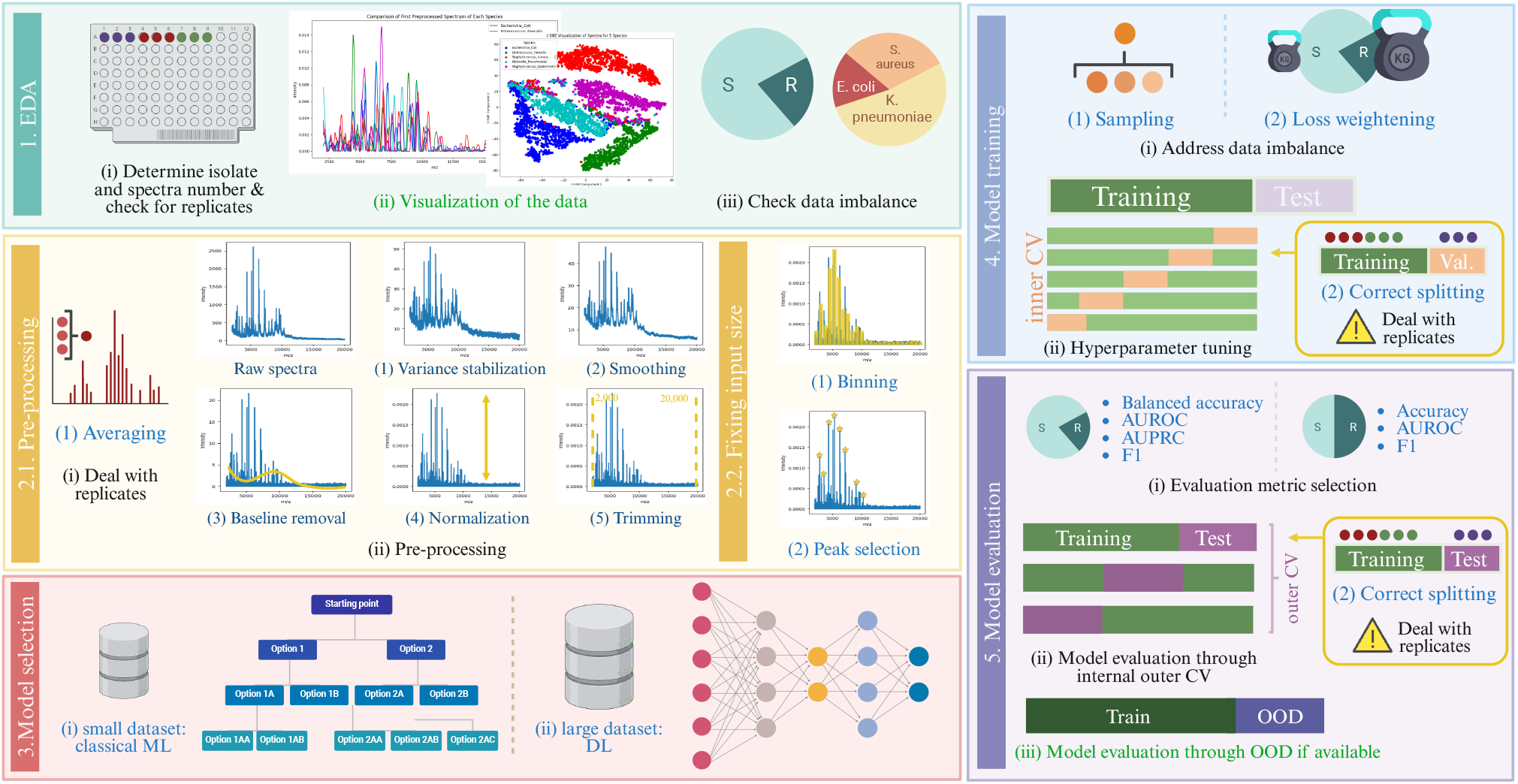
Overview of key steps in ML-based MALDI-TOF MS analysis, covering exploratory data analysis (EDA), preprocessing, model selection, training, and evaluation. The diagram highlights critical considerations such as data imbalance handling, hyperparameter tuning, and cross-validation (CV). Recommended steps are indicated in **black**, steps with multiple valid approaches are shown in blue, and optional steps are marked in green.

During the EDA, a comprehensive dataset analysis is crucial for making informed pre-processing and model selection decisions. One of the first aspects to consider is the number of unique bacterial isolates, as this directly influences model selection. Additionally, handling spectral replicates is essential, and there are two primary approaches: (1) either peak alignment and averaging spectra to create a single representation per isolate or (2) treating them as a multi-instance problem. The latter approach is preferable, as it maximizes data usage and improves reproducibility, but care must be taken to ensure that replicates remain within the same split in both inner and outer CV to prevent data leakage. Another key factor is identifying data imbalance, which has a direct impact on loss functions, and evaluation strategies. Finally, data visualization using unsupervised learning techniques, e.g. PCA or UHCA, can enhance interpretability and pattern recognition. However, these methods should be reserved for exploratory analysis rather than classification tasks, which require supervised learning models.

Pre-processing plays a fundamental role in standardizing datasets for better model learning. Although current SOTA practices lack consistency, we recommend several essential steps for raw spectral data: (i) variance stabilization using log2 or square root transformation, (ii) smoothing with a Savitzky-Golay filter, (iii) baseline removal with techniques such as the Statistics-sensitive Non-linear Iterative Peak-clipping algorithm (SNIP) or top-hat transformation, (iv) normalization of the intensities, and (v) trimming within the range of 2,000 to 20,000 Da to eliminate noise. After this, spectra can either be used in their original varying lengths or standardized. A straightforward approach is binning with a 3 Da bin size, converting spectra into 6,000-element vectors by summing intensities within each bin. Alternatively, PS techniques, e.g. based on RF, can be employed to retain the most relevant features. For more advanced implementations, models such as Transformers [114] can directly handle variable-length spectra.

Choosing the right model depends on the research question and the characteristics of the dataset. For small to medium-sized datasets, classical ML models such as RF and LGBM generally provide strong performance. Bayesian models, such as GPs [25] or FA [89], are particularly useful for combining multimodal data, as they generalize well and do not require an inner CV. In contrast, larger datasets allow for the use of DL models [21, 114], which are highly effective at capturing complex representations. Before model training, data imbalance must be addressed, which can be done through either (1) oversampling techniques like SMOTE [113, 114] or more targeted approaches [33], or by (2) incorporating imbalance-aware loss functions, such as weighted Log-Loss for logistic regression, weighted Hinge Loss for SVMs, or class-weight adjustments in RF models. Hyperparameter tuning is also a key step and should be performed using inner CV, calculating the mean and standard deviation of evaluation metrics to identify the most robust model. As in previous steps, spectral replicates must remain within the same split to prevent bias.

Model evaluation should follow training using outer CV, where the dataset is split into *k* folds (typically 3, 5, or 10), allowing the model to be trained and tested *k* times. Once the best-performing model is identified, it should be retrained using the entire internal dataset. To assess generalizability, an OOD evaluation should be performed using a completely independent dataset, ideally from a different machine or hospital. Evaluation metrics should address class imbalance, with balanced accuracy as a priority, complemented by AUROC, precision, recall, and AUPRC for a comprehensive performance assessment.

## Conclusions

The integration of ML into MALDI-TOF MS analysis has significantly advanced bacterial ID, AMR prediction, and strain TYP. However, several challenges remain, including data scarcity, insufficient transparency in reporting methodological details, non-standardized pre-processing workflows, inconsistencies in model evaluation strategies, and limited availability of open-source datasets and code. While classical ML models remain the predominant and often highly competitive approach, recent evidence suggests that increased model complexity does not inherently guarantee superior performance. Instead, performance gains appear strongly dependent on task complexity, dataset characteristics, and the ability to learn robust spectral representations. In this context, representation learning strategies, including transfer learning and self-supervised pre-training, emerge as key drivers of improvement, particularly for complex prediction scenarios. Despite these methodological advances, issues related to reproducibility, robustness, and generalizability continue to hinder reliable clinical deployment.

We hope this review not only provides a comprehensive overview of the current landscape but also serves as a guide toward more standardized and methodologically coherent practices in ML-based MALDI-TOF research. By promoting open science, transparent reporting, and rigorous external validation, we aim to support the development of robust, interpretable, and clinically relevant models. Furthermore, we seek to inspire future efforts to better align modelling strategies with clinical realities, foster interdisciplinary collaboration between data scientists and clinical experts, and ultimately drive broader clinical adoption of ML-enhanced MALDI-TOF MS technologies.

## Acknowledgements

This work was supported by the Ministry of Science, Innovation and Universities – Agencia Estatal de Investigación (MCIN/AEI/10.13039/501100011033) [grant number PID2023-146684NB-I00 to V.G.V.]; and the Community of Madrid [project TEC-2024/COM-89 to V.G.V.]. A.G.L. is supported by the European Union’s Horizon Europe programme through a Marie Sk-lodowska-Curie Actions Postdoctoral Fellowship (OUTBRAID, Grand Agreement No. 101278403). The work of C.S.S. was supported in part by the Comunidad de Madrid’s 2025 Cesar Nombela program under Grant 2025-T1/COM-36091.

## Data Availability

This article is a systematic review of previously published studies. No new datasets were generated or analyzed during the current study.

## Declaration of generative AI and AI-assisted technologies in the writing process

During the preparation of this work, the authors used ChatGPT (OpenAI) in order to enhance the clarity and quality of the manuscript, particularly for language refinement and consistency. After using this tool/service, the author(s) reviewed and edited the content as needed and take(s) full responsibility for the content of the publication.

## Footnotes

1 The search query was: ‘machine learning’ OR ‘classification’ OR ‘support vector machine’ OR ‘random forest’ OR ‘logistic regression’ OR ‘neural network’ OR ‘deep learning’ OR ‘transformer’ OR ‘multi-layer perceptron’ OR ‘embedding’ OR ‘clustering’ OR ‘unsupervised’ OR ‘self-supervised’ OR ‘semi-supervised’ OR ‘selfsupervised’ OR ‘semisupervised’ OR ‘bayesian’ OR ‘kernel’ AND ‘maldi-tof’.

2 To avoid confusion with citations, the number of studies is denoted as n=X.

